# Systemic endotoxemia induces integrated sickness physiology in female BALB/c mice

**DOI:** 10.64898/2026.08.22.746462

**Authors:** Pooja Kher, Bruna G. C. Lima, Cheyanne Woodrow, Ana C. Roginski, Leslie B. Hernandez, Anjali Wilson, Zeinab Tashi, Benjamin B. Bartelle, Esther B. Florsheim

## Abstract

Sickness is an organismal response to inflammation, yet its immune, metabolic, neural, and behavioral components are often studied separately and predominantly in male C57BL/6 mice. In this study, we characterized these responses to systemic lipopolysaccharide (LPS) in female BALB/c mice. Mice received intraperitoneal LPS at moderate concentrations and outcomes were assessed during the acute and resolving phases of endotoxemia. LPS caused rapid disappearance of resident peritoneal macrophages, followed by neutrophil accumulation and increased circulating TNF-α and IL-6. In the liver, LPS induced inflammatory, acute-phase, and anti-inflammatory transcripts while suppressing genes involved in lipid, cholesterol, and xenobiotic metabolism. Hepatic glutathione was reduced, whereas total superoxide dismutase activity was unchanged. These peripheral responses were followed by transient hypothermia, reduced food intake, and body weight loss. Regional brain mapping showed increased c-Fos labeling in the area postrema, nucleus of the solitary tract, external lateral parabrachial nucleus, paraventricular nucleus of the hypothalamus, and arcuate nucleus. In parallel, LPS selectively promoted IBA1-positive area in the median eminence and arcuate nucleus, whereas several other regions showed no changes, indicating that neuronal and microglial responses are regionally distinct. Behaviorally, LPS reduced locomotion and exploration, increased freezing, and increased forced-swim immobility. Changes in spatial exploration were most pronounced during the acute phase, whereas locomotor suppression and passive stress-coping persisted longer and varied in magnitude with the timing of inflammatory challenge. Together, these findings show that systemic LPS produces a coordinated sickness state in female BALB/c mice that links peripheral inflammation and hepatic metabolic and redox changes with region-specific neuronal and microglial responses, altered thermoregulation and feeding, and behavioral suppression.

## INTRODUCTION

Inflammation triggered by infection affects physiology far beyond the site where pathogens are detected. During acute illness, animals develop a recognizable set of physiological and behavioral changes, including altered body temperature, reduced food intake, lethargy, fatigue, disrupted sleep, and reduced grooming and social interest. These responses, often grouped under the term sickness behaviors, are familiar features of infection in animals (1–3). Although they are often experienced as symptoms, many sickness-associated responses are thought to reflect organized physiological programs that reduce competing demands and help animals cope with inflammatory stress (2,4–8). Yet, the contribution of individual sickness reactions to host defense and recovery remains incompletely understood. This distinction is clinically important because responses that are beneficial during acute infection may become harmful when excessive or prolonged, contributing to fatigue, cachexia, mood disturbance, and other complications of chronic inflammation (9–12). Understanding how immune activation is translated into coordinated changes in physiology and behavior therefore remains an important problem in physiology.

Systemic administration of lipopolysaccharide (LPS), a component of the outer membrane of Gram-negative bacteria, is widely used to model acute endotoxemia and sickness physiology in rodents and humans (13,14). Recognition of LPS by Toll-like receptor 4 initiates neutrophil recruitment and the production of cytokines such as interleukin (IL)-1β, IL-6, and tumor necrosis factor (TNF)-α, while also engaging hepatic acute-phase, metabolic, and redox programs (15–17). At the organismal level, LPS can produce fever or hypothermia, hypophagia, weight loss, reduced locomotion, and behavioral inhibition, with the nature and magnitude of these responses depending on dose and experimental context (1,18). Systemic inflammation is communicated to the brain through multiple routes, including circulating cytokines and lipid mediators, brain-vascular and circumventricular interfaces, and vagal afferent pathways (19–21). These signals recruit brainstem and hypothalamic regions involved in visceral sensing, thermoregulation, feeding, neuroendocrine responses, and behavioral state (20,22–24). In addition to altering neuronal activity, systemic LPS can induce local inflammatory reactions within the brain, including cytokine production and changes in astrocyte and microglial reactivity (25,26). Hence, the central response to peripheral inflammation includes both rapid neural signaling and regionally selective neuroimmune responses that may evolve over different time scales.

Despite extensive use of systemic LPS as a model of acute inflammation, many studies have examined immune, metabolic, neural, or behavioral outcomes in isolation, and much of the literature has focused on male rats or C57BL/6 mice (1,27). Comparatively less is known about how these responses are coordinated in female BALB/c mice, a strain widely used in immunological and allergic disease models (28–30). In this study, we characterized the integrated response to systemic LPS in female BALB/c mice across peripheral inflammation, liver transcriptional and redox responses, thermoregulation, feeding, neuronal recruitment, regional microglial IBA1 immunoreactivity, and behavior. By examining these responses within the same experimental framework, we sought to define how acute systemic inflammation reorganizes physiology across organ systems and to establish a foundation for mechanistic studies of sickness in this sex and genetic background.

## RESULTS

### Systemic LPS promotes prototypical inflammatory and hepatic acute-phase responses in female BALB/c mice

To establish an endotoxemia model in female BALB/c mice, we administered intraperitoneal injections of *Escherichia coli* lipopolysaccharide (LPS) at 0.5 or 1.0 mg/kg, non-septic doses previously used to induce nonlethal systemic inflammation and moderate sickness-associated responses in C57BL/6 or BALB/c mice (19,31–34). Inflammatory responses were assessed 6 and 24 h post-injection (Figure 1A). Manipulations were conducted starting at ZT14, reflecting the active phase of mice, a nocturnal species. Total number of cells within the peritoneal cavity nearly doubled at 24 h following LPS administration, whereas no significant increase was observed at 6 h (Figure 1B). Both LPS doses produced comparable cellular influx at 24 h. Flow cytometric analysis revealed that approximately 15% of infiltrating cells at 6 h and 50% of infiltrating cells at 24 h were neutrophils (Ly6G^+^F4/80^-^) (Figures 1C and S1A-B). Notably, although total cell numbers were unchanged at 6 h, LPS induced a dose-dependent reduction in resident macrophages (F4/80^hi^) accompanied by a reciprocal increase in neutrophil frequency (Figures 1D and S1C). This rapid macrophage loss is consistent with the classical “macrophage disappearance reaction”, originally described decades ago (35,36) and later linked to activation of omental B-1 B cell–mediated antibacterial defense (37,38). At 24 h, LPS also increased FcεRI⁺ cells within the peritoneal cavity (Figure S1D), which in this compartment are predominantly mast cells, as basophils are present at very low frequency (39). It is possible that these mast cells contribute to anti-bacterial defense, as previous study showed that the conditional depletion of mast cells increases mice susceptibility and lethality in a severe sepsis model (39).

**Figure 1.**
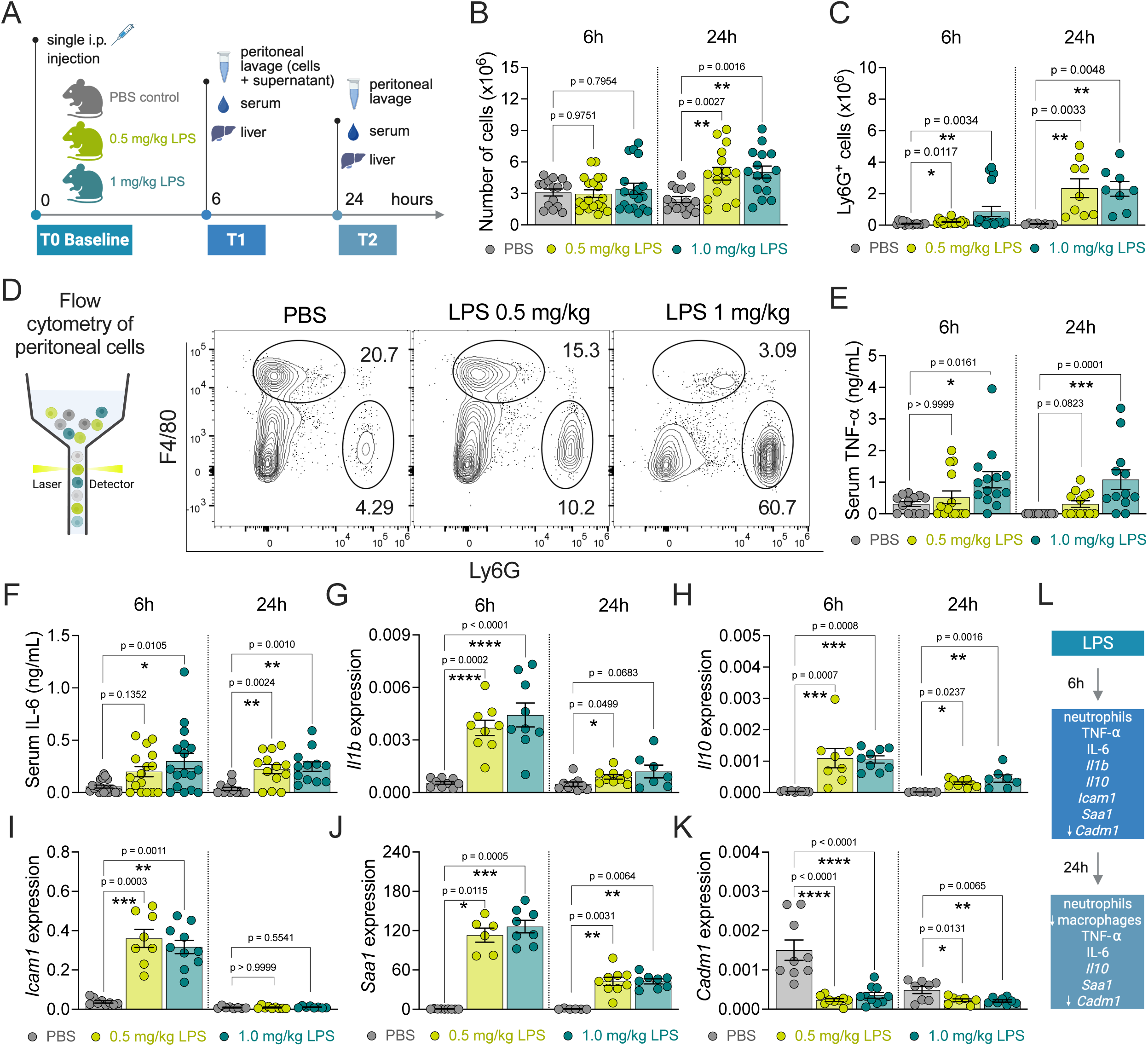
LPS treatment induces local and systemic inflammation in female BALB/c mice. **(A)** Study timeline and experimental groups. Peritoneal (cells and supernatant), serum, and liver samples were collected 6 or 24h post intraperitoneal injection of female BALB/c (7-8 w.o.) with bacterial lipopolysaccharide (LPS). Controls received PBS. **(B)** Total number of cells in the peritoneal cavity quantified with Trypan Blue. **(C)** Number of neutrophils (Ly6G^+^ cells, gated from viable and singlets). **(D)** Schematic of flow cytometry for immunophenotyping of peritoneal cells (left). Frequency of neutrophils (Ly6G^+^) and macrophages (F4/80^hi^) in the peritoneal cavity 6h post injection (right). **(E-F)** Serum concentrations of TNF-α and IL-6 quantified by ELISA. **(G-K)** Liver expression of interleukin 1 β (*Il1b)*, interleukin 10 (*Il10)*, intercellular adhesion molecule 1 (*Icam1)*, serum amyloid A1 (*Saa1)*, and cell adhesion molecule 1 (*Cadm1)* relative to *Rpl13a* quantified by RT-qPCR. (**L**) Summary of findings. Graphs show mean±s.e.m. *p<0.05, **p<0.01, ***p<0.001, ****p<0.0001. Kruskal–Wallis test with Dunn’s post-test (E, F, C-6h, G-6h, H, I) or one-way ANOVA test with Dunnett’s post-test (B, C-24h, G-24h, J, K). Each panel is representative of at least two independent experiments. A, D, and L, created with BioRender.

Systemically, circulating tumor necrosis factor (TNF)-α and interleukin-6 (IL-6) were significantly elevated at 24 h, whereas at 6 h only the higher LPS dose induced robust increases (Figures 1E–F). Together, these findings demonstrate that systemic LPS administration in female BALB/c mice elicits a coordinated innate inflammatory response characterized by neutrophil recruitment, macrophage disappearance, mast-cell accumulation, and systemic cytokine production.

Because activation of the hepatic acute-phase response is a hallmark of systemic inflammation, we next examined liver transcriptional programs following LPS exposure. Robust induction of pro- and anti-inflammatory and acute-phase transcripts was evident as early as 6 h, including interleukin-1β (*Il1b*), tumor necrosis factor-α (*Tnfa*), interleukin-6 (*Il6*), interleukin-10 (*Il10*), intercellular adhesion molecule-1 (*Icam1*), serum amyloid A1 (*Saa1*), arachidonate 5-lipoxygenase (*Alox5*), and histidine decarboxylase (*Hdc*) (Figures 1G–J and S1E–H). At 24 h, most transcripts remained elevated, albeit at lower magnitudes, whereas *Icam1*, *Il6*, and *Alox5* returned toward baseline relative to the housekeeping gene *Rpl13a*. In contrast, leukotriene C4 synthase (*Ltc4s*) was not induced at 6 h and was reduced at 24 h in a dose-dependent manner (Figure S1I). Together with increased *Alox5* expression, this pattern suggests reduced hepatic capacity for cysteinyl-leukotriene synthesis and a potential shift toward LTB_4_ production (40,41). As expected, transcripts of prostaglandin E synthase 2 (*Ptges2*), a GSH-dependent enzyme constitutively expressed in the liver and involved in the generation of a metabolite of arachidonic acid (PGE2), were mildly upregulated by i.p. LPS (Figure S1J). Expression of cell adhesion molecule 1 (*Cadm1*) was acutely and persistently reduced (Figure 1K), whereas vascular endothelial growth factor A (*Vegfa*) was downregulated as early as 6 h (Figure S1K). Together with a modest reduction in transforming growth factor-β1 (*Tgfb1*) expression (Figure S1L), these changes are consistent with temporary suppression of hepatic tissue-repair and angiogenic programs. In contrast, hepatic expression of interferon-β1 (*Ifnb1*) and chymase 1 (*Cma1*) was not consistently altered by LPS (Figures S1M–N).

Collectively, these data establish that systemic LPS exposure in female BALB/c mice induces a robust inflammatory state encompassing local innate immune cell mobilization, systemic cytokine release, and activation of hepatic acute-phase and lipid-mediator transcriptional programs (Figures 1L and S1O). Because such inflammatory responses are known to engage organism-level regulatory circuits governing thermoregulation, metabolism, and energy balance, we next investigated whether this endotoxemic state is followed by coordinated physiological alterations characteristic of sickness.

### Systemic LPS elicits coordinated sickness physiology and hepatic stress responses in female BALB/c

We next evaluated the dynamic physiological consequences of systemic LPS exposure in female BALB/c mice (Figure 2A). Non-lethal LPS administration produced a transient hypothermic response, with rectal temperature reaching its nadir approximately 2 h post-injection and returning to baseline by 6 h (Figure 2B, left). Quantification of the maximal temperature change (Δ temperature) revealed that LPS quickly reduced body temperature in a dose-dependent manner, by approximately 2°C in the highest concentration of 1 mg/kg (Figure 2B, right). Systemic LPS exposure also resulted in an expected body-weight loss, pronounced at 6 h and maintained at 24 h (Figure 2C), followed by reduced food intake at both time points (Figure 2D). These results indicate coordinated metabolic suppression during acute inflammation.

**Figure 2.**
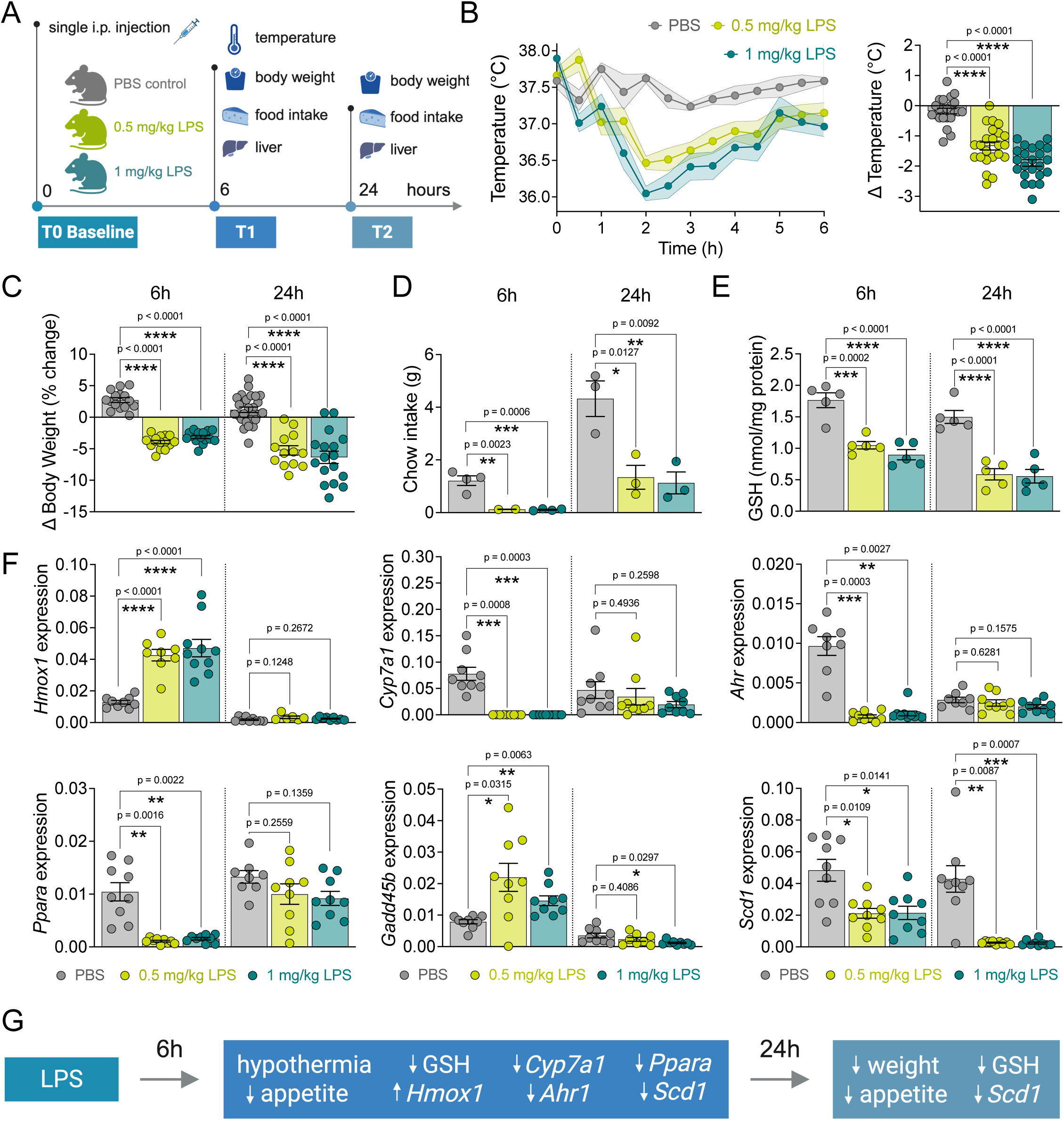
LPS induces coordinated physiological changes in female BALB/c mice. (**A**) Study timeline and experimental groups. Read-outs included the analysis of rectal temperature, body weight, food intake, and liver samples post intraperitoneal injection of female BALB/c (7-8 w.o.) with LPS. Controls received PBS. (**B**) Time course of rectal temperature post injections (left) and maximal temperature change calculated as 2-hour time point temperature minus initial (right). (**C**) Percentage of change in body weight at 6 h and 24 h post-injection. (**D**) Chow intake (g per mouse per cage). (**E**) Liver concentration of reduced glutathione (GSH) in nmol/mg protein. (**F**) Hepatic expression of heme oxygenase 1 (*Hmox1*), cholesterol 7 α-hydrolase (*Cyp7a1*), growth arrest and DNA damage inducible β (*Gadd45b*), aryl hydrocarbon receptor (*Ahr*), peroxisome proliferator-activated receptor α (*Ppara*), and stearoyl-CoA desaturase-1 (*Scd1*), normalized to housekeeping gene *Rpl13a* quantified by RT-qPCR. Graphs show mean±s.e.m. *p<0.05, **p<0.01, ***p<0.001, ****p<0.0001. Group differences were tested using ordinary one-way ANOVA (B, C and E) when assumptions were met and Kruskal– Wallis (F) for non-normally distributed data. Panels A and G were created with BioRender.

Given that endotoxemia is associated with oxidative stress (42–44), we next assessed hepatic redox balance. Liver glutathione (GSH) levels were reduced by nearly 50% as early as 6 h post-LPS and remained reduced 24 h later (Figure 2E), consistent with the development of acute oxidative stress. We then examined superoxide dismutase pathways to characterize antioxidant regulation. Expression of superoxide dismutase 1 (*Sod1*) was transiently reduced at 6 h but mostly normalized by 24 h (Figure S2A). Despite the transient reduction in *Sod1* transcript levels, total superoxide dismutase (SOD) enzymatic activity was not altered at any LPS dose or time point examined (Figure S2B), indicating partial preservation of antioxidant enzymatic capacity during the acute inflammatory response. Next, we examined hepatic expression of 5-oxoprolinase (*Oplah*), which is essential for glutathione recycling, and found that LPS did not change its expression (Figure S2C).

To further define hepatic responses to LPS-induced systemic inflammation, we analyzed transcriptional programs associated with antioxidant defense, metabolic regulation, and cellular stress signaling. LPS induced a transient increase in heme oxygenase 1 (*Hmox1*) at 6 h that returned to baseline by 24 h (Figure 2F). In contrast, genes involved in cholesterol and xenobiotic metabolism and lipid homeostasis—including cholesterol 7α-hydroxylase (*Cyp7a1*), aryl hydrocarbon receptor (*Ahr*), and peroxisome proliferator-activated receptor α (*Ppara*)—were robustly suppressed at 6 h and normalized by 24 h (Figure 2F). The stress-response regulator growth arrest and DNA-damage-inducible β (*Gadd45b*) showed a modest increase at 6 h followed by a slight decrease at 24 h following the higher LPS dose. Finally, the lipogenic enzyme stearoyl-CoA desaturase 1 (*Scd1*) exhibited a mild reduction at 6 h that became more pronounced at 24 h, indicating sustained disruption of hepatic lipid metabolism.

Together, these findings demonstrate that systemic LPS exposure in female BALB/c mice triggers a coordinated physiological syndrome characterized by transient hypothermia, weight loss associated with reduced appetite, acute hepatic oxidative stress with selective modulation of antioxidant pathways, and temporally structured transcriptional reprogramming of metabolic and stress-response programs (Figure 2G).

### Systemic LPS engages brainstem-hypothalamic circuits and elicits region-specific microglial responses

To characterize central responses to peripheral inflammation, we quantified expression of the immediate-early gene Fos as a proxy for neuronal activation following LPS administration (45) (Figure 3A). For subsequent brain and behavioral analyses, we selected a single LPS dose of 1 mg/kg because it induced robust systemic cytokine production and neutrophil recruitment (Figure 1), whereas most other inflammatory and physiological responses were comparable between doses.

**Figure 3.**
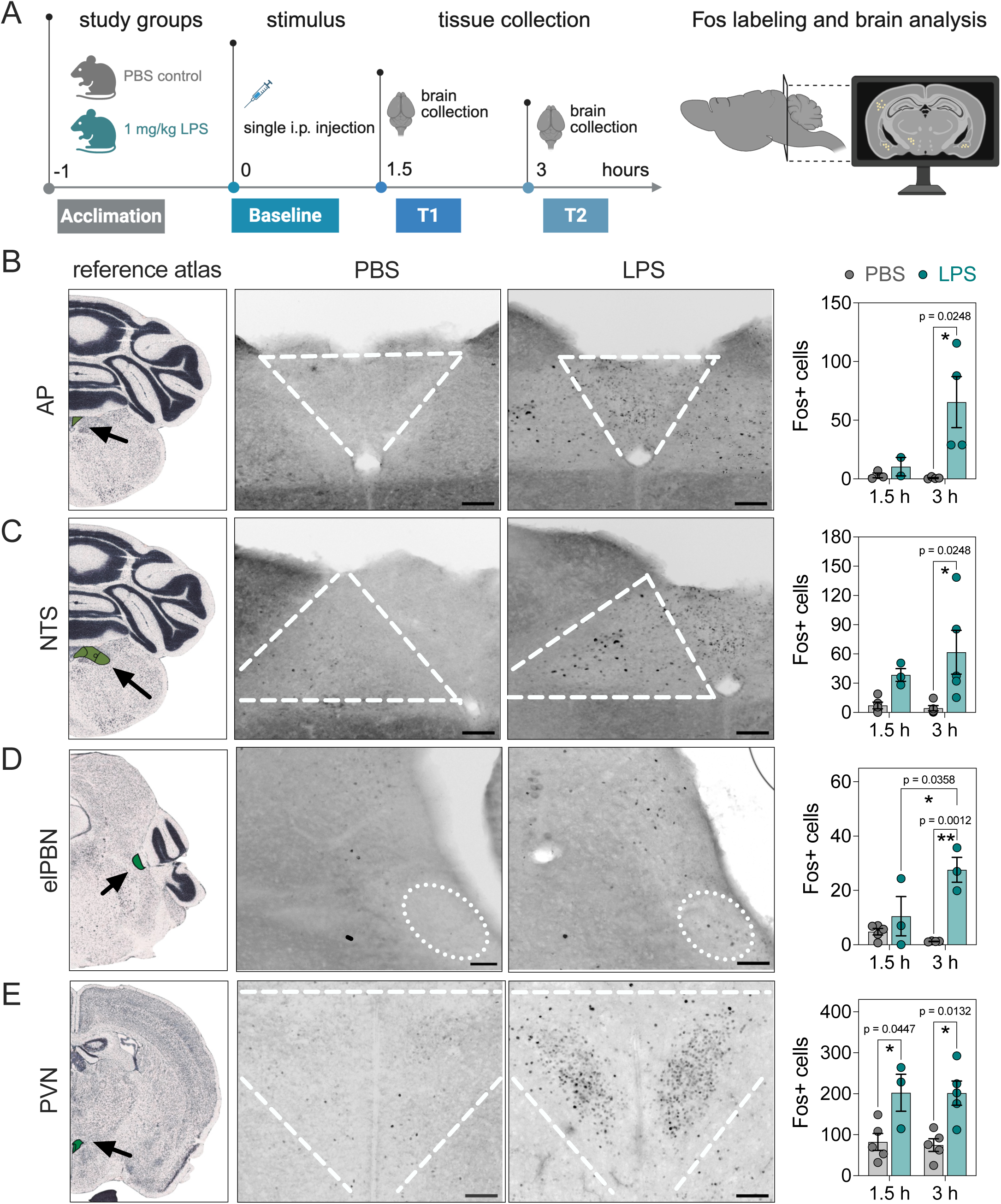
LPS-induced brain activation in female BALB/c mice. (**A**) Schematic protocol to assess neuronal activation following i.p. LPS (1 mg/kg) in female BALB/c mice. Brains were collected 1.5 h or 3 h post LPS or PBS (controls), sliced coronally, and imaged for Fos labeling and quantification. (**B-E**) From left to right: images from the Allen Mouse Brain Atlas, followed by coronal brain sections from PBS- and LPS-treated mice. Number of Fos+ cells per region per time point. Abbreviations: AP: area postrema; NTS: *nucleus tractus solitarius*; elPBN: external lateral portion of the parabrachial nucleus; PVN: paraventricular nucleus of the hypothalamus. Graphs show mean±s.e.m. Two-way ANOVA was used to compare group differences. *p<0.05, **p<0.01. Scale = 170μm. Panel A was created with BioRender.

Systemic LPS is known to recruit brainstem and hypothalamic regions involved in interoception and homeostatic regulation (22,46,47). Consistent with these studies, regional mapping in female BALB/c mice revealed stimulus-evoked c-Fos labeling in several of these areas. Prominent responses were observed in the area postrema (AP) and nucleus of the solitary tract (NTS) (Figures 3B-C), which together integrate circulating and visceral inflammatory signals. In the pons, LPS also increased the number of c-Fos^+^ cells within the parabrachial nucleus (PBN), particularly within its external lateral subregion (Figure 3D), a region implicated in processing aversive and malaise-associated signals (48). In the hypothalamus, the paraventricular nucleus (PVN) showed robust c-Fos induction at 3 h, with increased labelling already detectable at 90 min (Figure 3E). In contrast, minimal Fos labeling in these regions was observed in vehicle-treated controls.

LPS did not significantly alter neuronal activation in the posterior paraventricular nucleus of the thalamus (pPVT) or central amygdala (CeA) at the time points examined (Figure S3A-B).

In the dentate gyrus (DG), c-Fos labeling was reduced 3 h after LPS administration (Figure S3C), indicating decreased neuronal recruitment during acute endotoxemia. Conversely, we found increased neuronal activation in the arcuate nucleus of the hypothalamus (ARC) of LPS-treated mice compared with controls (Figure S3D), consistent with recruitment of a homeostatic region involved in the regulation of feeding and metabolism.

We next determined whether systemic LPS was followed by regional changes in microglial IBA1 immunoreactivity (Figure 5A-F). The AP showed a higher IBA1-positive area compared to neighboring regions like the NTS, under control conditions. This basal activation is consistent with previous studies of circumventricular organs (CVOs) and was not further increased by LPS (Figure 5G). The lower baseline IBA1 immunoreactivity of the NTS likewise did not show a significant change following LPS treatment (Figure 5G). Microglia of the midbrain CVO, the median eminence (ME), did not have the same basal activation as the AP. In contrast, LPS exposure significantly increased IBA-1-positive area exclusively in the ME and ARC, whereas no significant differences were detected in the DG, ventromedial hypothalamus (VMH), and corpus callosum (CC) (Figure 5G). Therefore, the microglial response to systemic LPS was regionally selective, with most prominent changes occurring in the ME and neighboring ARC.

Together, these findings identify distinct regional neuronal and microglial responses to systemic LPS. Neuronal recruitment was prominent across brainstem and hypothalamic regions involved in visceral sensing and homeostatic regulation, whereas increased microglial IBA1 immunoreactivity was restricted primarily to the ME and ARC. These patterns suggest that neuronal and microglial responses to systemic inflammation are spatially differentiated rather than uniformly coupled across LPS-responsive brain regions (Figure 5H).

### Acute LPS induces coordinated sickness-associated behavioral states

We next characterized the behavioral consequences of systemic LPS exposure in female BALB/c mice (Figures 4A–B). Locomotor activity and spatial exploration were assessed using the open field test, whereas behavioral persistence during an acute stressor was evaluated using the forced swim test. Animals were recorded using automated video-tracking software, which generated spatial trajectory maps (track plots) to evaluate arena exploration. Under control conditions, mice exhibit moderate thigmotaxis, a natural tendency to remain near the walls while intermittently exploring the center of a novel arena. Increased thigmotaxis, reflected by greater occupancy of peripheral or corner zones and reduced center exploration, is commonly interpreted as an anxiety-like behavioral response (49,50). To align behavioral testing with the inflammatory and physiological measurements described above, and to better reflect the nocturnal activity pattern of *Mus musculus*, mice were injected at ZT14 during the dark phase.

**Figure 4.**
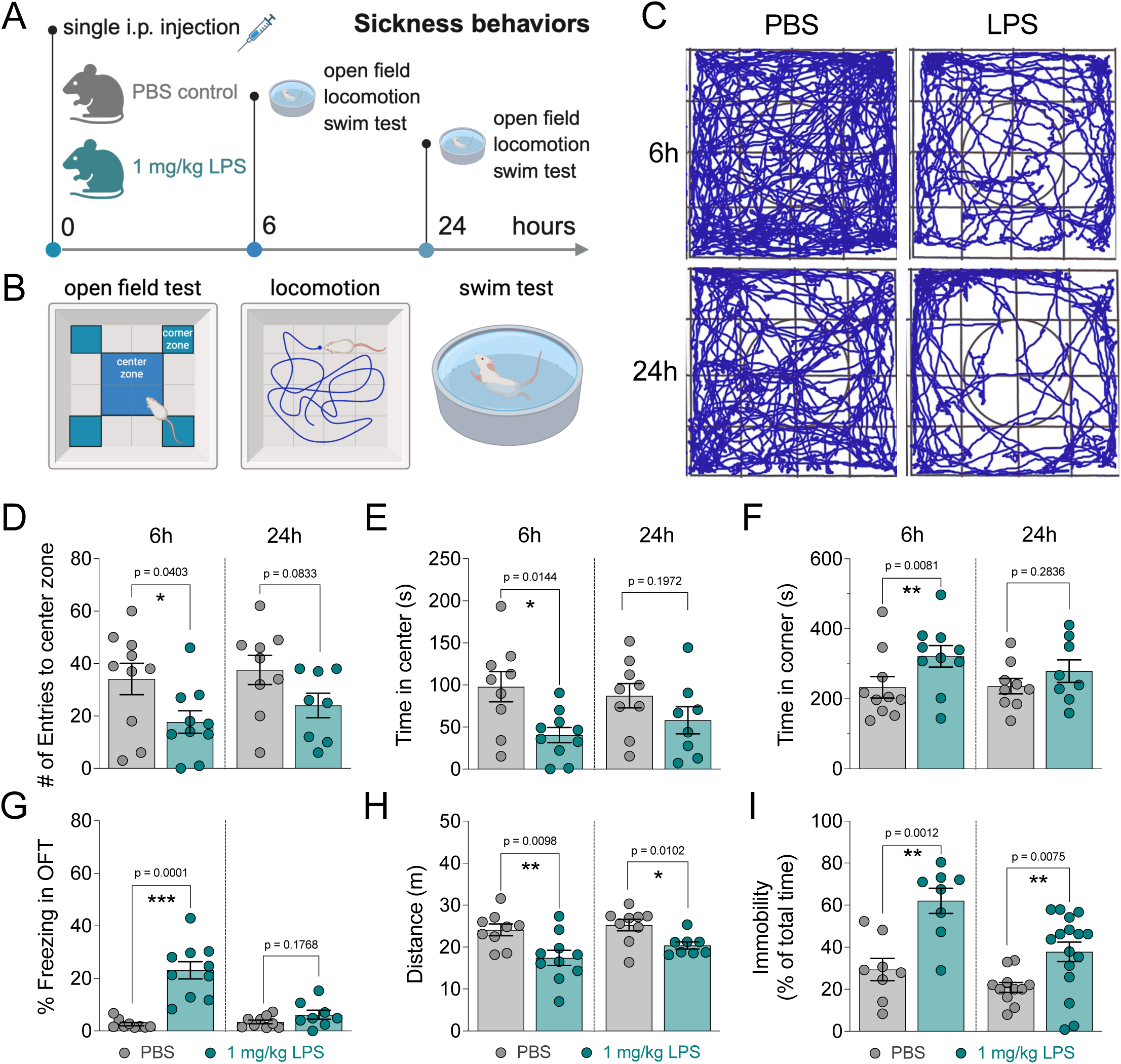
Peripheral LPS induces sickness behaviors in female BALB/c mice. (**A**) Study timeline and experimental groups. Sickness behaviors were analyzed at 6 or 24h post intraperitoneal injection of female BALB/c with 1 mg/kg LPS. Controls received PBS. (**B**) Behavioral assays included the open field test, locomotion, and forced swim test. Mice were acclimated to the behavioral room 1 h prior to testing under controlled red-light conditions. **C**) Representative track plots during a 10-min session in the open field arena. (**D**) Number of entries to the center zone during OFT. (**E**) Time spent in the center zone and (**F**) time spent in the corner zone. (**G**) Percentage of time freezing during the OFT, defined as immobility bouts lasting longer than 7s. (**H**) Total distance traveled in the OFT. (**I**) Percentage of immobility during the forced swim task. Graphs show mean±s.e.m. Data at 6 h were pooled from two independent experiments. Abbreviation: OFT, open field test. Group differences per time point were tested using unpaired student t test (D-I). *p<0.05, **p<0.01, ***p<0.001. Panels A and B were created with BioRender.

**Figure 5.**
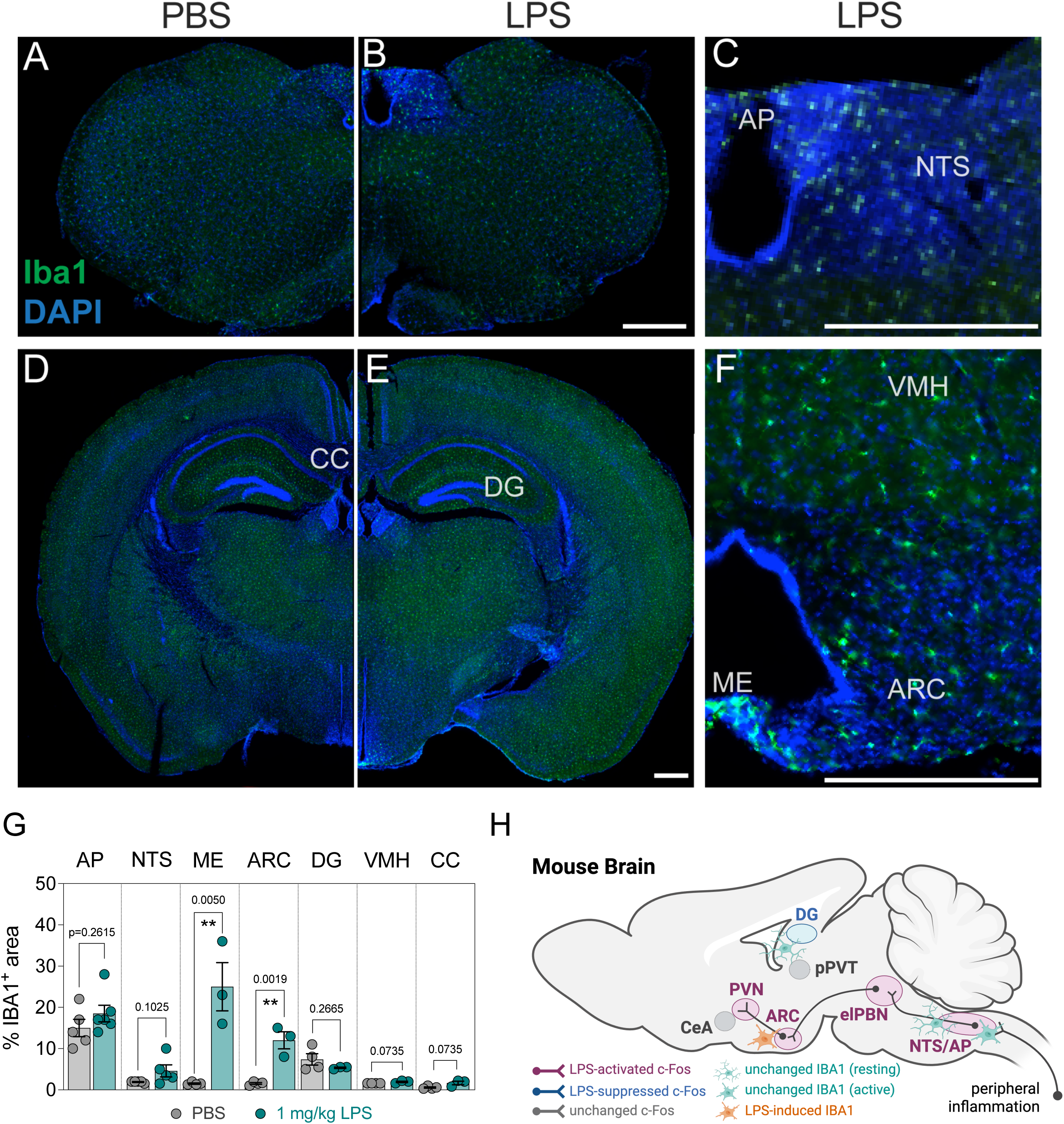
Microglial activation by LPS. Female BALB/c mice were i.p. injected with 1mg/kg LPS. Controls received PBS alone. Brains were collected 24h later upon perfusion. (**A**) Immunofluorescence (IF) of the mouse brainstem for PBS group. (**B**) Brainstem post LPS stimulation. (**C**) IF of the AP and NTS. (**D-E**) Mouse midbrain for PBS and LPS-treated mice, respectively. (**F**) IF of ME, ARC, and VMH. (**G**) Pixel quantification for brainstem (AP and NTS) and midbrain (ME, ARC, DG, VMH, CC) regions. (**H**) Schematic summary of neuronal and microglial responses to systemic LPS in the female BALB/c brain. Scale bar= 400 µm. Graphs show mean±s.e.m. Abbreviations: AP, area postrema; NTS, nucleus of the solitary tract; ME, median eminence; ARC, arcuate; VMH, ventromedial hypothalamus; DG, dentate gyrus; CC, corpus callosum. Group differences per time point were tested using unpaired student t test. **p<0.01. H, created with Biorender.

Representative track plots showed reduced exploratory trajectories after LPS treatment at both 6 and 24 h compared with vehicle-treated controls (Figure 4C). Deeper analysis of open field behavior revealed that LPS-treated mice entered the center zone less frequently as early as 6 h post-injection (Figure 4D). Spatial analysis further showed that LPS reduced time spent in the center zone and increased time spent in corner regions at 6 h (Figures 4E-F). This redistribution of arena occupancy was no longer evident at 24 h, suggesting that LPS-induced changes in exploratory patterns are transient. Animals treated with LPS also exhibited increased freezing behavior at 6 h (Figure 4G), consistent with enhanced behavioral inhibition during acute systemic inflammation. Quantitative analysis confirmed that LPS-treated mice traveled less distance during the 10-min open field task compared with controls at 6 h post-injection, and this reduction in locomotor activity persisted, although more modestly, at 24 h (Figure 4H). In contrast, passive stress-coping behavior occurred as early as 6 h post challenge and remained strongly affected at the later time point, as LPS exposure significantly increased the percentage of immobile time in the forced swim test relative to vehicle-treated controls (Figure 4I).

To determine whether circadian phase influences LPS-induced behavioral responses, mice were injected at ZT5 during the inactive light phase, and behavioral outputs were analyzed 24 h later. Consistent with the dark-phase experiments, LPS did not alter preference for the center or corner zones of the open field arena at this time point (Figure S4A-B). LPS-treated mice showed a slight reduction in locomotion compared with controls (Figure S4C), comparable to the residual locomotor impairment observed when animals were injected during the dark phase (Figure 4H). However, immobility in the forced swim test was more pronounced when LPS was administered during the light phase (Figure S4D). These findings suggest that LPS administration during the inactive phase produces a more persistent passive stress-coping state and locomotor impairment, without inducing lasting changes in center avoidance or exploratory zone preference. Therefore, some components of LPS-induced sickness behavior are influenced by circadian phase.

Altogether, these results indicate that systemic LPS exposure in female BALB/c mice induces a coordinated sickness behavioral syndrome characterized by suppressed locomotion, transient alterations in exploratory behavior, increased behavioral inhibition, and robust changes in passive stress-coping behavior. These behavioral outputs temporally coincide with activation of brainstem and hypothalamic circuits engaged during systemic inflammation.

## DISCUSSION

During acute inflammation, physiological processes, including behavior, are reorganized as part of an integrated response that may help organisms to cope with inflammatory stress and restore homeostasis. Here, we show that systemic LPS exposure in female BALB/c mice produces a coordinated sickness state spanning peritoneal and systemic inflammation, hepatic acute-phase and redox responses, altered thermoregulation and feeding, selective neuronal and microglial activation in the brainstem and hypothalamus, and sickness-associated behavioral changes. Previous studies have often examined these components separately or focused predominantly on male animals and C57BL/6 mice. Our findings therefore provide an integrated reference for studying sickness physiology in female BALB/c mice and a foundation for future comparisons across sex, genetic background, and experimental context/Female BALB/c mice mounted a temporally organized innate immune response to LPS. Early disappearance of resident peritoneal macrophages occurred alongside increasing neutrophil frequency, before the later increase in the total number of cells and neutrophil migration. This response was accompanied by circulating TNF-α and IL-6 and by a broad hepatic inflammatory and acute-phase program. The liver response included not only pro-inflammatory transcripts such as *Il1b*, *Tnf*, and *Il6*, but also *Il10* and histidine decarboxylase (*Hdc*), suggesting that counter-regulatory mechanisms are engaged alongside inflammatory amplification. LPS has been previously reported to induce HDC in the liver (51). Histamine, in turn, can suppress LPS-induced TNF-α through H2 receptors and has been implicated in the regulation of LPS-induced liver inflammation (52–54). Because we measured *Hdc* expression rather than histamine concentrations or cellular sources, these findings do not establish a histaminergic mechanism but are consistent with increased histamine-synthetic capacity. Similarly, increased *Alox5* together with reduced *Ltc4s* expression suggests a transcriptional shift away from cysteinyl-leukotriene synthesis and a potential shift toward LTB4 production, although direct lipid-mediator measurements will be required to confirm this interpretation. The sustained reduction in hepatic *Cadm1* expression was also notable, although the function and cellular source of CADM1 in acute hepatic inflammation remain poorly defined and will require further investigation.

LPS also induced substantial hepatic metabolic and redox reprogramming. Hepatic GSH was reduced, whereas total SOD activity remained unchanged, indicating selective disruption of redox balance rather than a generalized loss of antioxidant enzymatic function. This is consistent with evidence linking LPS-induced oxidative changes in the liver to symptoms of sickness (55). These changes were accompanied by transient induction of *Hmox1* and suppression of transcripts involved in bile acid synthesis, xenobiotic sensing, fatty acid regulation, and lipogenesis, including *Cyp7a1*, *Ahr*, *Ppara*, and *Scd1*. Previous work in male BALB/c mice similarly found that LPS induced hepatic inflammatory cytokines and heme oxygenase-1 while suppressing PPAR expression (56), and TLR–TRIF signaling can directly repress the transcription of stearoyl-coenzyme A (CoA) desaturase 1 (*Scd1*), the rate-limiting enzyme for lipogenesis, in hepatocytes (57). Recent work showed that heme released during tissue injury triggers oxidative stress and an NRF2-dependent HMOX1 detoxification program in macrophages (58). Although heme was not measured here, this finding provides another potential context for the transient hepatic induction of *Hmox1* after LPS. Sustained LPS-induced inflammation has similarly been shown to suppress CYP7A1 and redirect hepatic mevalonate-pathway metabolism, with consequences for systemic glucose homeostasis (59). Although we did not measure glucose production or metabolic pathway flux, the early reduction in *Cyp7a1* suggests that inflammation-driven reorganization of hepatic cholesterol and bile acid metabolism begins rapidly. These findings position the liver as an active participant in the redistribution of metabolic resources during sickness. In the absence of histopathology or circulating markers of hepatocellular damage, however, they are best interpreted as metabolic and oxidative stress-associated reprogramming rather than exacerbated liver injury.

The physiological response was characterized by transient hypothermia together with longer lasting hypophagia and weight loss. Thermoregulatory responses to LPS depend on dose, route, ambient temperature, sex, strain, and the timing and method of measurement (18,60). Lower doses frequently induce fever, whereas stronger inflammatory challenges, particularly under sub-thermoneutral conditions, can produce hypothermia. Previous comparisons showed that BALB/c mice develop reduced body temperature after intraperitoneal LPS, although the magnitude varies across sex and strain (61). In our model, temperature reached its nadir approximately 2 h after LPS and returned toward baseline by 6 h, whereas reduced feeding and weight loss persisted through 24 h, indicating that individual components of sickness recover on distinct time scales.

Hypothermia and hypophagia may also represent energy-conserving components of disease tolerance rather than nonspecific physiological failure (62). During severe LPS- and *E. coli*–induced systemic inflammation, allowing animals to develop spontaneous hypothermia reduced organ dysfunction and overall mortality compared with experimentally imposing fever (63). Hypothermia and hypometabolism can preserve host function without necessarily reducing inflammatory burden or pathogen load (64,65). Reduced feeding can similarly engage fasting-associated metabolic programs that support essential physiology during bacterial inflammation, including hepatic FGF21-dependent maintenance of thermoregulation and cardiovascular function (8,66). Loss of appetite promoted by LPS has also been shown to take priority even when eating behavior is stimulated (22). Thus, the hypothermia, hypophagia, and hepatic metabolic changes observed here may represent a coordinated energy-management strategy. Their adaptive value in this nonlethal model remains uncertain, however, because we did not manipulate temperature or feeding or assess their effects on tissue damage and survival.

The pattern of neuronal recruitment observed here was centered on regions positioned to detect and integrate peripheral inflammatory signals. LPS increased c-Fos labeling in the area postrema (AP), nucleus of the solitary tract (NTS), external lateral parabrachial nucleus, paraventricular nucleus of the hypothalamus, and arcuate nucleus (ARC). This pattern overlaps with classical LPS-responsive regions whose recruitment depends partly on prostaglandin signaling (19) and with causal evidence that ADCYAP1-positive AP/NTS neurons regulate feeding, drinking, locomotion, and temperature during LPS-induced sickness (22). Distinct hypothalamic circuits also contribute to opposing thermoregulatory responses: ventromedial preoptic area neurons coordinate fever and warmth seeking after LPS challenge (23), whereas a region at the ventral border of the dorsomedial hypothalamus is required for cold seeking during high-dose LPS-induced hypothermia (24). Although we did not manipulate these circuits, recruitment of a broader brainstem-hypothalamic network is consistent with an organized hypothermic sickness response. The absence of significant central amygdala (CeA) activation contrasts with reports of CeA recruitment after LPS in male BALB/c mice (33). Notably, oral allergen challenge robustly increases CeA activation in sensitized female BALB/c mice (30), arguing against a generalized inability of this sext-strain combination to engage this region and suggesting that CeA recruitment may depend on the nature of the peripheral immune stimulus. Reduced Fos labeling in the dentate gyrus (DG), in turn, indicates diminished neuronal recruitment during acute endotoxemia but does not by itself imply impaired memory.

Systemic LPS can also elicit inflammatory responses within the brain. At comparable doses, peripheral LPS has been reported to induce brain cytokine expression, astrocyte reactivity, and increased immunoreactivity of the microglial marker ionized calcium-binding adapter molecule 1 (IBA1) in the hippocampal DG (25,26). In our study, LPS increased IBA-1-positive area selectively in the median eminence (ME) and ARC, whereas no significant changes were detected in the AP, NTS, DG, ventromedial hypothalamus, or corpus callosum. Thus, regional changes in microglial IBA1 immunoreactivity did not simply parallel neuronal c-Fos responses. The preferential response in the ME and ARC is relevant given their proximity to circulating signals and their roles in hypothalamic control of metabolism and feeding. Whether the increased IBA1-positive area reflects changes in microglial morphology, density, or activation state, and whether these responses contribute to LPS-induced physiological or behavioral changes, remains to be determined.

The behavioral effects of LPS were consistent with a coordinated sickness-associated state. At 6 h, LPS reduced locomotion and center exploration while increasing corner occupancy and freezing. By 24 h, spatial preference had largely normalized, whereas locomotor suppression and forced-swim immobility remained elevated. Previous work in male BALB/c mice similarly showed reductions in locomotion, feeding, social interaction, and novel-object exploration after LPS administration (33), and BALB/c mice of both sexes can show prolonged locomotor effects following systemic inflammation by LPS (31). These assays nevertheless require cautious interpretation during acute illness. Fatigue, malaise, hypophagia, altered motivation, and reduced locomotor capacity can all affect open field and forced-swim performances. The transient reduction in center exploration may include an anxiety-like component, but here it occurred within a broader state of hypoactivity and behavioral inhibition. Likewise, increased forced-swim immobility is better interpreted as reduced behavioral persistence or passive stress-coping than as evidence of a primary depressive state. This distinction is consistent with experimental and translational studies showing that sickness-related hypoactivity complicates conventional affective interpretations and that anxiety- and depression-related outcomes do not map straightforwardly between rodent and human LPS models (14,67,68).

Time of day also influenced the duration or magnitude of some behavioral responses. The main experiments were initiated at ZT14 to align the challenge with the active phase of this nocturnal species. When LPS was administered at ZT5 and behavior was assessed 24 h later during the light phase, center and corner occupancy were unchanged, whereas locomotor suppression remained detectable and forced-swim immobility was more pronounced. These findings suggest that exploration, general activity, and passive stress-coping do not necessarily recover in parallel and may be influenced differentially by the temporal context of inflammation and their respective role in physiological adaptation. Because challenge and testing times were shifted together, the current design cannot distinguish effects of challenge time from effects of testing time or establish direct circadian clock regulation.

More broadly, the responses observed here support the view that sickness is not a single uniform output of inflammation. A recent study comparing bacterial, viral, allergic, parasitic, and intestinal inflammatory challenges across organismal physiology and behavior, brain-wide neuronal activity, and cell-type-specific transcriptional responses found that distinct immune insults generated different multiscale sickness states (47). In this context, LPS induces one particularly broad sickness program, but its expression may still vary with sex, genetic background, dose, and time of day. Our findings extend this emerging framework by defining how the LPS-induced state is organized across peripheral inflammation, liver physiology, brain activity, and behavior in female BALB/c mice. In this framework, sickness is not simply a collection of symptoms but an organized and context-dependent state that promote disease tolerance and reallocates organismal resources during inflammatory stress.

### Limitations, caveats, and open questions

We provide an integrated description of LPS-induced sickness physiology in female BALB/c mice, but future studies will be required to establish causal relationships among inflammation, brain activation, and behavior. Fos and IBA1 labeling identifies recruited regions and morphology, respectively, but cannot determine whether they are necessary or sufficient for individual physiological or behavioral responses. We did not measure brain cytokines or astrocyte reactivity, circulating or hepatic histamine, leukotriene concentrations, whole-body energy expenditure, or markers of hepatocellular injury. Rectal temperature measurements also provide discrete estimates and may not capture complex thermal dynamics detectable by continuous telemetry. In addition, LPS is a controlled model of bacterial endotoxemia and does not reproduce pathogen replication or host–pathogen interactions. In summary, systemic LPS exposure in female BALB/c mice induces an integrated sickness state spanning innate immune activation, hepatic inflammatory and metabolic reprogramming, redox disruption, altered thermoregulation and feeding, selective brainstem and hypothalamic recruitment, and sickness-associated behavioral suppression. This model provides a foundation for studying how peripheral inflammatory signals are integrated across organ systems to generate and resolve whole-animal responses to acute inflammation.

## METHODS

### Mice

BALB/c mice (strain code 028) were initially purchased from the Charles River Laboratories and maintained in our laboratory. Female adult mice at 7 to 12 weeks old were used throughout the study. Mice were maintained at the Arizona State University animal facilities under specific pathogen-free conditions. Mice were euthanized by CO_2_ asphyxia. All protocols were reviewed and approved by the Arizona State University Institutional Animal Care and Use Committee. Mice were group-housed (up to 5 animals per cage) in standard polycarbonate cages placed on a suspended-cage rack with water and food *ad libitum* on a 12 h dark/light cycle (lights on at ZT0, 7 pm). Room temperatures were maintained around 22°C, with humidity between 30-40%.

### Key Resources Table

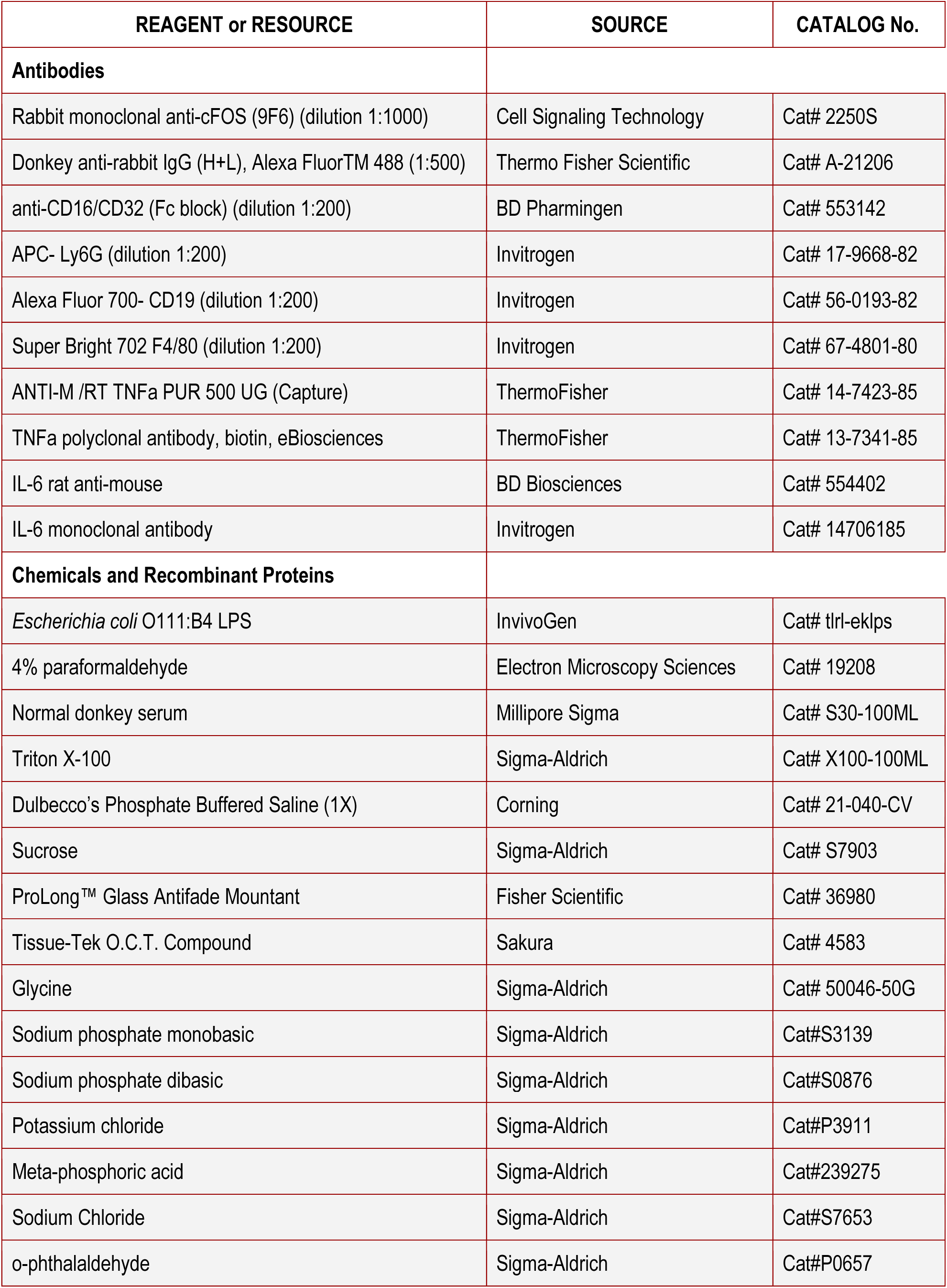
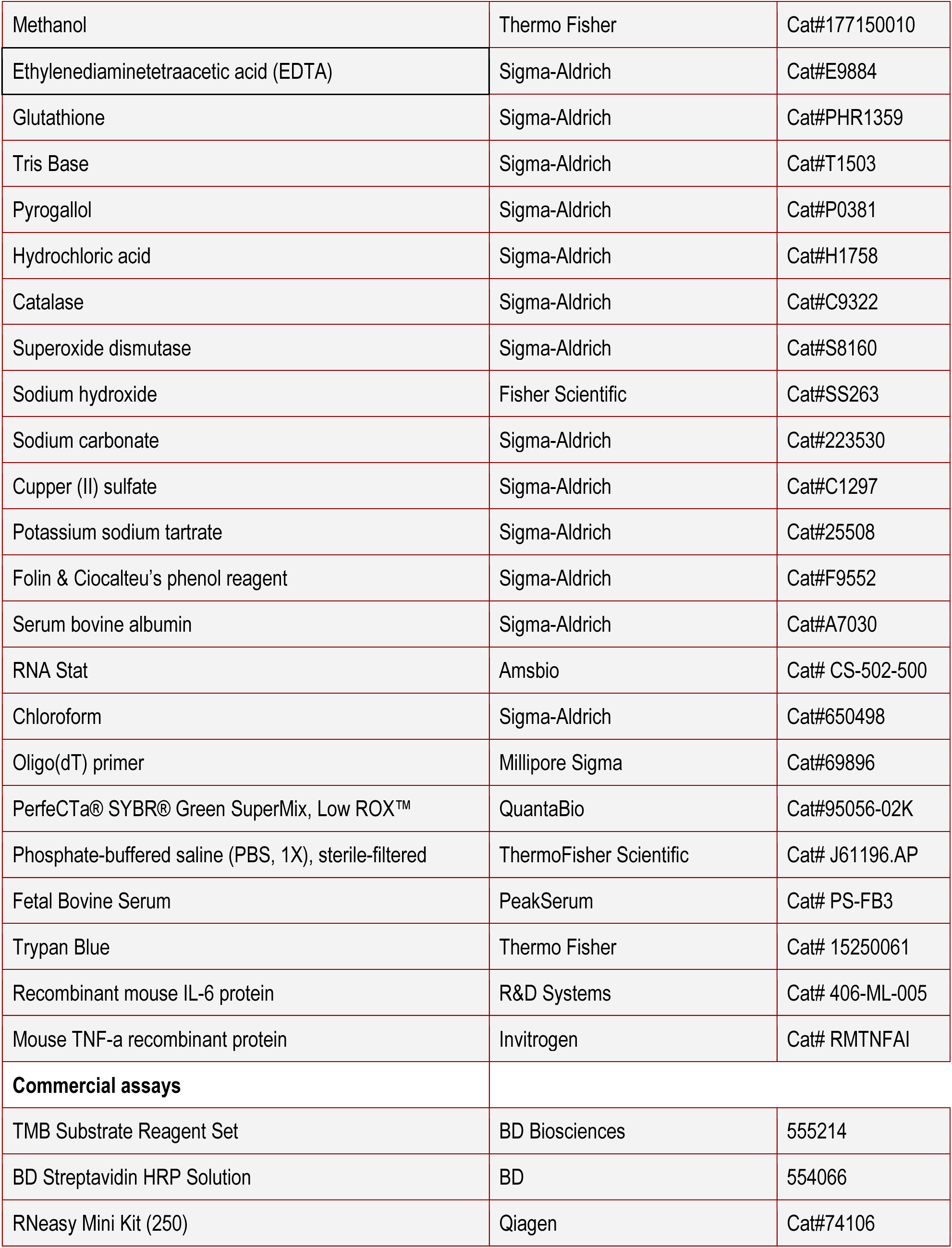
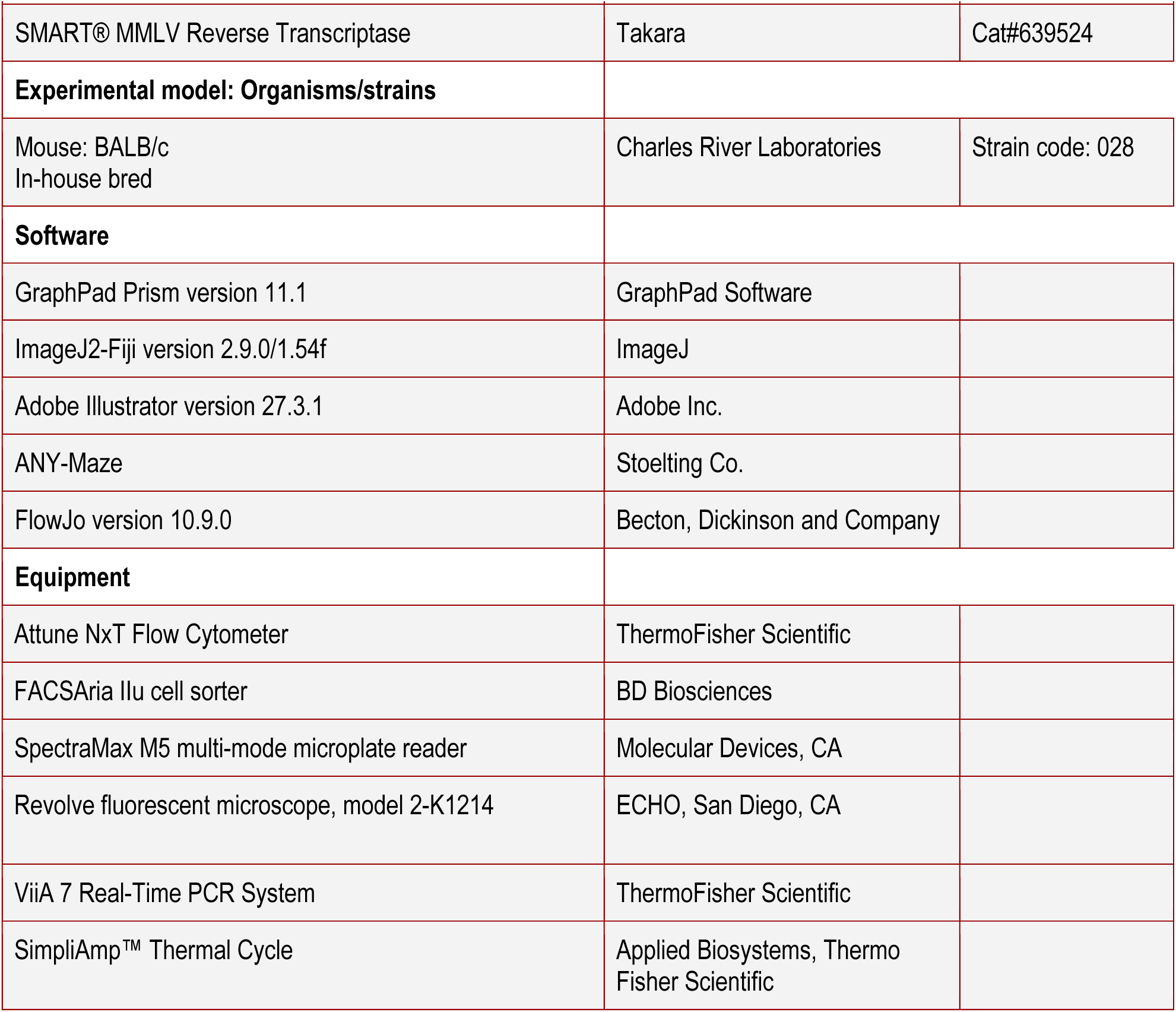

### Systemic inflammation model

To induce acute systemic inflammation, mice were intraperitoneally injected with 0.5 mg/kg or 1 mg/kg of lipopolysaccharides (LPS from *E. coli*; Invivogen) in 0.2 mL of sterile PBS. Control mice were i.p. injected with the same volume of PBS. Six or twenty-four hours later, mice were euthanized and serum, peritoneal lavage, liver or brain tissues were collected for cytokine ELISA, flow cytometry, mRNA expression analysis, immunofluorescence, and analysis of oxidative stress. The peritoneal lavage was collected by injecting and recovering 4 mL of cold sterile PBS into the abdominal cavity. The total number of peritoneal cells was quantified using a 0.4% solution of Trypan Blue (Thermo Fisher, 15250061) and a Neubauer chamber hemocytometer. Serum was terminally collected via cardiac puncture immediately after euthanasia.

### Flow cytometry

Peritoneal cells were collected via peritoneal lavage utilizing cold PBS. Cell suspensions were centrifuged at 570 g for 5 min and washed twice with ice-cold PBS (Thermo Fisher Scientific), and cell counts were then collected. Peritoneal cell suspensions were divided into separate aliquots and stained in parallel with distinct antibody panels. Pelleted cells were stained with Zombie Yellow-Viability dye (BioLegend, 77168, 1:1000) for 15 min at RT. Cells were then washed twice with ice-cold FACS buffer made with PBS (Thermo Fisher Scientific) supplemented with 2% FBS (Fetal Bovine Serum, Peak Serum) and stained with fluorochrome-coupled antibodies APC-Ly6G (Invitrogen, 17-9668-82, clone: 1A8-Ly6g 1:200), Alexa Fluor 700-CD19 (Invitrogen, 56-0193-82, clone: eBio1D3 1:200), Super Bright 702-F4/80 (Invitrogen, 67-4801-80, clone: BM8 1:200), PerCP-eflour 710-CD3 (Invitrogen, 46-0032-82, clone: 17A2, 1:200), FITC-MHCII (Invitrogen, 11-5321-85, clone: M5/114.15.2, 1:200), PE-FcER1 (Invitrogen, 12-5898-81, clone: MAR-1, 1:200), and anti-CD16/32 (Fc block; BD Pharmingen, 553142, 1:200) in 2% FBS in PBS for 30 min at 4°C protected from light. Cell suspensions were then centrifuged and washed twice with PBS supplemented with 2% FBS (PS-FB3, Peak Serum, Inc), followed by incubation with PBS and 2% PFA (paraformaldehyde, 20% w/v aq. soln., methanol free, ThermoFisher Scientific, 047340.9M) for 25 min at 4°C protected from light. Cells were acquired using the Attune NxT Flow Cytometer (ThermoFisher Scientific, B2R3V6Y3) with 405, 488, 561, and 640 nm excitation lasers and the FACSAria IIu cell sorter (BD Biosciences) with 405, 488, 561, and 640 nm excitation lasers. Analysis was carried out using FlowJo™ Software (Becton, Dickinson, and Company). The gating strategy was drawn from negative controls, first gating to isolate cells from debris, followed by single cells from doublets, and then viable cells (Zombie Yellow-negative). Neutrophils were defined as Zombie Yellow–negative, CD19-negative, F4/80-negative, Ly6G-positive.

### Cytokine ELISA

Serum and peritoneal lavage supernatants were assayed for levels of TNF-α and IL-6 by sandwich enzyme-linked immunosorbent assay (ELISA). ELISA-grade plates (Thermo Fisher, 442404) were coated with 2 μg/mL of anti-mouse TNF-α (eBioscience, 13-7423-85) or anti-mouse IL-6 (eBioscience, 14-7061-85) in 0.1M sodium carbonate buffer (pH 9.5) overnight at 4°C. Plates were blocked with 1% bovine serum albumin (Sigma-Aldrich, A7030) at room temperature (RT) for 1 h. Standards and serum from control and LPS-injected mice was diluted to 1:2.5, then incubated at RT for 2 h. Purified mouse TNF-α (Invitrogen, 29-8321-65) or IL-6 (R&D Systems, DY406-05) was used as a standard curve, with the highest concentration being 10ng/mL (TNF-α) or 4 ng/mL (IL-6 followed by two-fold dilutions. Cytokines were detected with biotin-conjugated anti-TNF-α (eBioscience, 13-7341-85) or anti-IL-6 (BD Biosciences, 554402) at 1 μg/mL for 1h followed by a1:1000-fold dilution of HRP-conjugated streptavidin (BD Biosciences, 554066) for 30 min at RT. Plates were then developed in TMB substrate reagent (BD Biosciences, 555214) in the dark at RT. Reactions were stopped with 2.5N sulfuric acid (Ricca, UN2796) at the appearance of a two-fold dilution on the standard curve. Absorbance at 450nm was read immediately using a plate reader (Varioskan Lux, Thermo Scientific, Waltham, MA). Between each step, plates were washed 3-7 times with 0.05% Tween-20 (Sigma-Aldrich, P1379) in PBS.

### RNA isolation and qPCR

For tissue RNA extraction, approximately 30 mg of liver tissue was harvested into RNA STAT-60 RNA isolation reagent (Amsbio, CS-502-500) and disrupted by bead homogenization (Omni, INC), then flash frozen on dry ice. RNA was extracted using the RNeasy Mini Kit (Qiagen, 74106) following the manufactures instructions. Briefly, 0.2 mL of chloroform was added to the 1mL of RNA-STAT 60 homogenized tissues, samples were mixed for 30 seconds, kept on ice for 5 min, and centrifuged at 12,000 g for 10 min at 4°C. Next, 0.35 mL of 70% ethanol was added to the same amount aqueous phase (RNA), and then was loaded onto Qiagen RNeasy column, the remainder of the manufacturer’s protocol was followed, including the on-column DNase digestion step. The final RNA pellet was eluted with 50 μL of nuclease-free water (Sigma). Total RNA was reverse transcribed using an oligo(dT) primer and Moloney murine leukemia virus reverse transcriptase (MMLV RT, TakaraBio). cDNA was analyzed by quantitative PCR amplification using Power SYBR™ Green PCR Master Mix (Life Technologies, 4367660) on a ViiA 7 Real-Time PCR System (Life TechnologiesTM, Applied Biosystems). Primers were designed to amplify mRNA-specific sequences, and analysis of the melt-curve confirmed the amplification of single products. Relative expression was normalized to ribosomal protein L13a (RPL13a). primers used are provided below.

### Primers Table

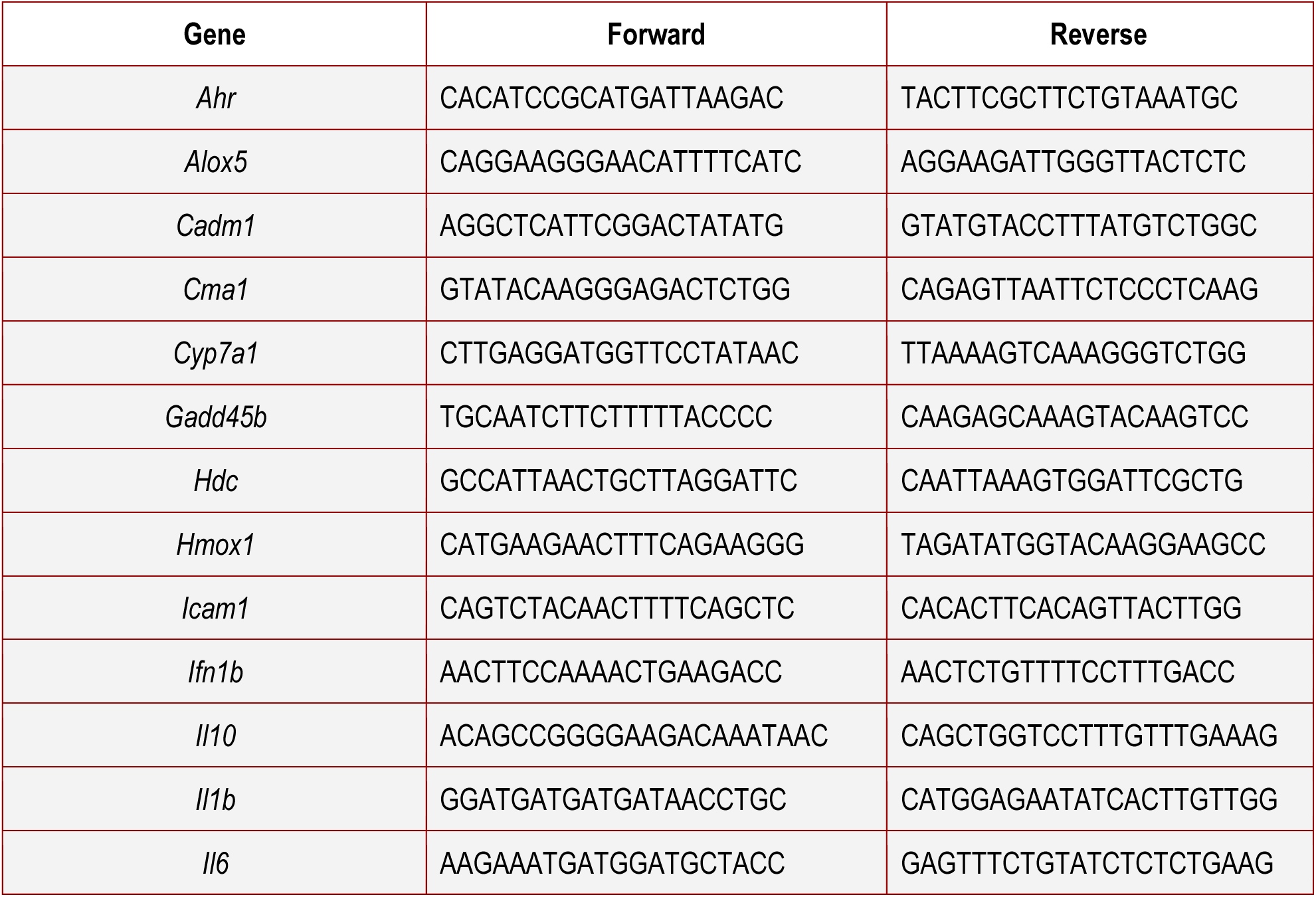
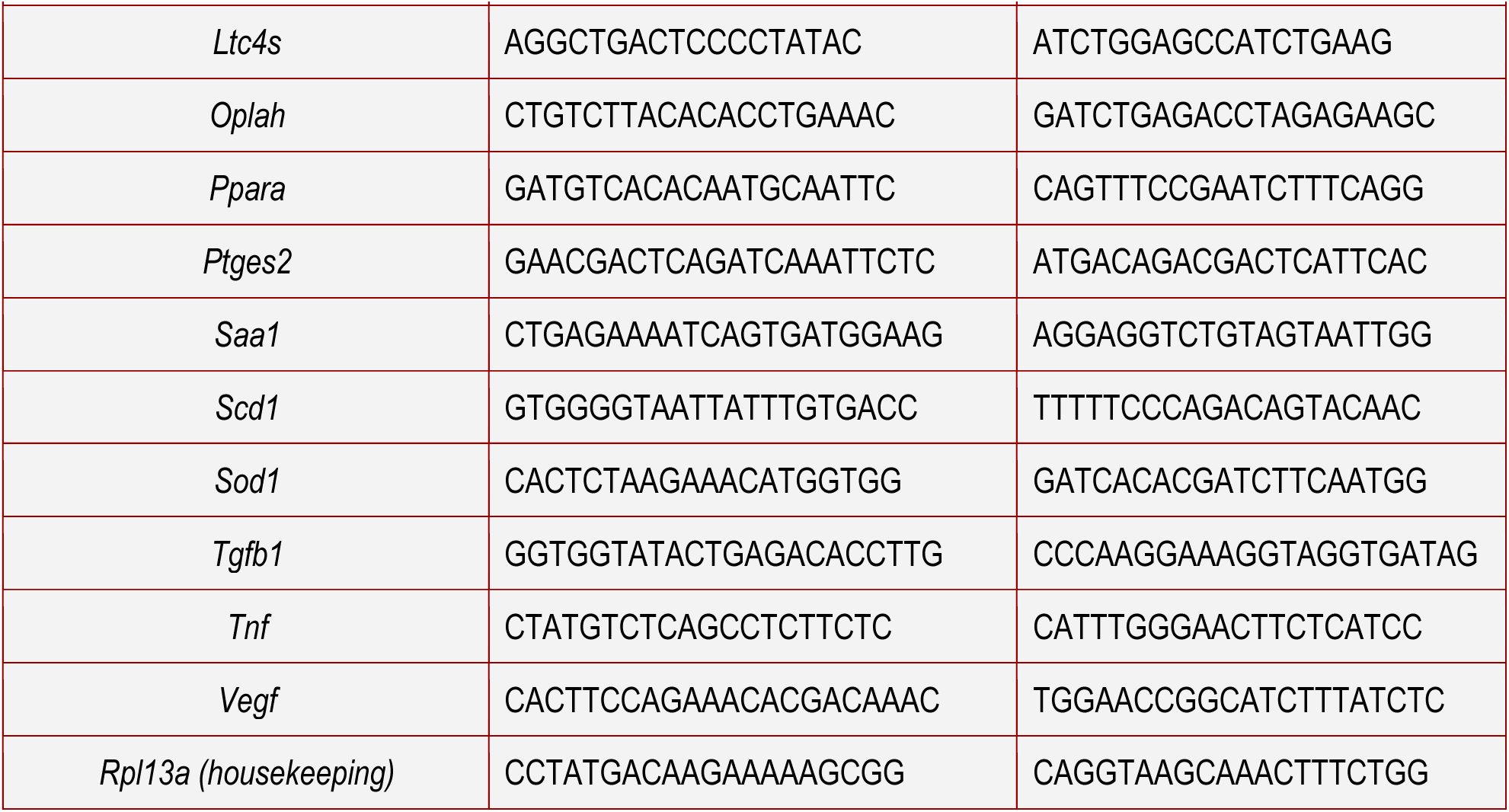

### Physiological measurements

#### Body weight

Mice were weighed immediately before injection (baseline) and at 6 h and 24 h after intraperitoneal injection.

#### Feeding

Food intake (in grams, g) was measured by weighing the food pellets offered per cage before and after the experiment (6 h and 24 h after injection). The final value refers to the weight difference between final and initial measurements divided by the number of mice per cage.

#### Temperature

A rectal probe was used to record body temperature every 30 min time up to 6 h post injections. Baseline temperature was recorded right before i.p. injections.

### Oxidative stress measurements

#### Glutathione

GSH concentrations were determined as previously described by Browne and Armstrong (69) with some adaptations. Briefly, 150 μL of liver supernatant was deproteinized with 2% metaphosphoric acid (1:1) and centrifuged for 10 min at 7000 g at 4°C. Later, 30 μL of deproteinized supernatant was added to a medium containing 185 μL of 100 mM sodium phosphate buffer, pH 8.0, containing 5 mM EDTA, and 15 mL of o-phthaldialdehyde (1 mg/mL in methanol) and incubated in the dark for 15 min at RT. Fluorescence was measured on a SpectraMax M5 plate reader (Molecular Devices, Sunnyvale, CA, USA), using excitation and emission wavelengths of 350 and 420 nm, respectively. A calibration curve was prepared using a standard GSH solution (0.001 – 1 mM), and results were expressed as nmol GSH/mg protein.

#### Superoxide dismutase

SOD activity was evaluated according to Marklund et al. (70) and is based on the capacity of pyrogallol to autoxidize, a process highly dependent on superoxide anion, which is the substrate for SOD. The inhibition of the autoxidation of pyrogallol occurs in the presence of SOD and, therefore, is proportional to the activity of the SOD present in homogenates. The reaction medium contained 50 mM Tris buffer, 1 mM ethylenediaminetetraacetic acid, pH 8.2, 80 U/mL catalase, 0.38 mM pyrogallol, and approximately 1 μg of protein, and the absorbance was read at 420 nm on a SpectraMax M5 plate reader (Molecular Devices, Sunnyvale, CA, USA). A calibration curve was constructed with purified SOD as standard to calculate the activity of SOD in the samples. The specific activity was calculated as U/mg protein.

For both assays, protein concentration was measured by Lowry et al. (71) method, using bovine serum albumin as a standard.

### Brain analysis

#### Brain collection and sectioning

For brain collections, mice were deeply anesthetized with 4% isoflurane. Mice were then perfused transcardially with PBS followed by 4% paraformaldehyde (pH 7.4). Once collected and dissected, brains were maintained in 4% PFA for 48 h at 4°C. Next, brains were washed with PBS and transferred to a 30% sucrose solution in PBS and kept in this solution until they sank. Brains were then covered with Tissue-Tek OCT, frozen and cut into 40 μm (from area postrema, AP, to anterior nucleus tractus solitarius, NTS) and 60 μm (from posterior NTS to prefrontal cortex) coronal sections using a cryostat (Leica CM1860, Leica Biosystems). Slices were stored in PBS at 4°C.

#### Neuronal activation

Brain slices were permeabilized for 30 min (PBS containing 0.3% triton X-100) at RT. Slices were then incubated in a blocking solution for 1 h (PBS containing 0.3% triton X-100 and 5% normal donkey serum) at RT. Slices were incubated with rabbit anti-cFos antibody (Cell Signaling Technology, 1:1000), overnight at 4°C. Next, slices were incubated with the secondary antibody donkey anti-rabbit IgG (H+L) Alexa Fluor 488 (Thermo Fisher Scientific, 1:500) for 2 h at RT. Finally, brains slices were mounted on slides and covered with a coverslip using Prolong Glass Antifade Reagent as the mounting media and stored at 4°C. Brain sections were imaged at a 10x magnification with a fluorescent microscope (Revolve, model 2-K1214, Echo, San Diego, CA). Fos positive neurons were quantified using the ImageJ2 software (ImageJ, version 2.9.0/1.54f) and expressed as the mean number of activated neurons within each area for at least 3 slices for each animal.

#### Microglia analysis

Brain sections were blocked with goat blocking serum (10% goat serum, 0.3% BSA, 0.1% Triton-X). Samples were incubated on a shaker at 80 rpm, 4°C overnight. On day 2, primary antibodies were added at 1:500 in goat blocking serum (chicken anti-IBA1 AB318302), and samples were kept on the shaker at 80 rpm at 4°C overnight. On day 3, samples were washed with PBS, followed by 3 times washing for a minimum of 10 min with PBS + 0.1% Triton-X, on a rocker. Secondary antibody (chicken AlexaFluor 488, Jackson Immuno Research 703-545-155) at 1:500 in goat blocking serum was added, and samples were kept on a shaker at 80 rpm at 4°C overnight. On day 4, samples were washed with PBS + 0.1% Triton-X for 10 min on the rocker, followed by 1:1000 diluted DAPI (ThermoFisher Cat#62248), and kept on the rocker for 15 min at RT. Finally, samples were washed 3 times with PBS on the rocker for at least 10 min each time, then placed on a charged glass slide. We added mounting media (Invitrogen ProLong Glass Antifade Mountant Cat#P36984), gently put the coverslip, and let the samples dry in the dark at RT until imaging (described above).

### Behavioral assays

A different batch of animals was used for each of the different timepoints. All behavioral experiments were performed under red light and with white noise (61 dB) unless mentioned otherwise.

#### Open Field Test (OFT)

Mice were acclimated for at least 1 h prior to behavioral assays. The effects of LPS on locomotor activity and anxiety-like behavior were assessed following a standard procedure as previously published (49) using an Open Field Chamber (18 in x 18 in x 12 in), whose floor was divided into 4 corners and a circle center. To abolish any olfactory cues, the chamber was wiped with Virex prior to use and between tests. Once each mouse was placed in the center of the arena, the test consisted of 2 min of habituation to reduce novel environmental stress, followed by 15 min to roam freely while being recorded with a Logitech® Webcam camera placed above the apparatus. Parameters such as speed, movement, immobility, and location in the arena were tracked and analyzed with the ANY-maze software connected to the camera.

#### Forced Swim Test (FST)

After exposure to OFT, mice were then subjected to the FST for a duration of 6 min. The FST tests depressive-like behavior by placing the mouse in a small, confined pool of water as previously reported (72,73). The apparatus consisted of a 5 L glass cylinder (25.6 cm H x 18.8 cm D) filled up to half with room temperature water (25°C). Each mouse was placed into the cylinder, and activity was monitored for 6 min by a Logitech® Webcam camera mounted placed in front of the apparatus. Mice were euthanized at the end of the behavioral protocols. Immobility time within the last 4 min of the test was hand-scored and analyzed by two blinded, independent researchers to this study.

### Quantification and statistical analysis

For all experiments, Shapiro-Wilk and Kolmogorov-Smirnov tests were used for normality test, α = 0.05, followed by either an unpaired t-test or Mann-Whitney test (two-group comparison) or by either an ordinary one-way ANOVA using Tukey’s multiple comparison test or Kruskal-Wallis using Dunn’s multiple comparison test to evaluate significance between three and more groups. The results were considered significant when p<0.05. Outliers were identified using ROUT’s method in GraphPad Prism at Q = 1%. All data were expressed as means ± s.e.m.

## ACKNOWLEDGMENTS

We would like to thank all current and former members of the Florsheim lab for helpful discussions. We thank the Department of Animal Care and Technologies team for technical assistance, and Biodesign Institute employees for their continuous assistance. Thanks to Zuri Sullivan for friendly reviewing the manuscript and providing critical feedback. Work in the Florsheim lab was supported by the Food Allergy Science Initiative (2022-FASI-0001, 2023-FASI-AS23310), the Hypothesis Fund (G11391-300), and the Arizona Department of Health (2024-022). Schematics were created with BioRender.com.

## Author contributions

P.K., B.C.L., C.W., B.B.B., and E.B.F. designed the study, analyzed the data, and wrote the manuscript with input from the other co-authors. P.K. performed behavioral experiments with the L.B.H. and A.W. assistance. A.C.R. performed the oxidative stress measurements and Z.T. performed the microglia staining. E.B.F. supervised the research.

## Declaration of generative AI and AI-assisted technologies

During preparation of this manuscript, the authors used ChatGPT (OpenAI) to improve conciseness, grammar, and readability. All suggested revisions were reviewed and edited by the authors, who take full responsibility for the final content of the manuscript.

## SUPPLEMENTAL FIGURE LEGENDS

**Supp. Figure S1 | LPS treatment induces local and systemic changes in immune cells profile and gene expression in female BALB/c mice.** (**A**) Gating strategy used to determine the frequencies of innate immune cells in the peritoneal cavity of a high dose LPS-treated mouse by flow cytometry. (**B**) Percentage of Ly6G+ cells (neutrophils) in the peritoneal cavity of PBS- and LPS-treated mice at 6h and 24h post injection. (**C**) Number of F4/80^+^ peritoneal cells 6h post LPS. (**D**) Number of FceRI^+^ peritoneal cells 24h post LPS. (**E-N**) Liver expression of tumor necrosis factor α (*Tnf)*, interleukin 6 (*Il6)*, arachidonate 5-lipoxygenase (*Alox5)*, histidine decarboxylase (*Hdc)*, leukotriene c4 synthase (*Ltc4s)*, prostaglandin E synthase 2 *(Ptges2),* vascular endothelial growth factor (*Vegf),* transforming growth factor β1 (*Tgfb1),* interferon β1 *(Ifnb1), and* chymase 1 (*Cma1)* relative to housekeeping gene *Rpl13a* quantified by RT-qPCR. (J-N) Panels show expression of liver transcripts at 6h post injections. (**O**) Summary of findings. Kruskal–Wallis test with Dunn’s post-test (B-6h, C-24h, D, E, F-24h, G, H-6h, I-24h, J, K, L, N) or One-way ANOVA test with Dunnett’s post-test (B-24h, H-24h, I-6h, M) was used. Data are presented as mean±s.e.m. *p<0.05, **p<0.01, ***p<0.001, ****p<0.0001. A and O were created with BioRender.

**Supp. Figure S2 | Hepatic antioxidant responses following LPS challenge. (A)** Hepatic expression of superoxide dismutase 1 (*Sod1*) at 6 h and 24 h after intraperitoneal injection of PBS, 0.5 or 1.0 mg/kg LPS. **(B)** Superoxide dismutase (SOD) activity (U/mg protein) in the liver. **(C)** Hepatic expression of 5-oxoprolinase (*Oplah*) at 6 h and 24 h relative to housekeeping gene *Rpl13a* quantified by RT-qPCR. Points represent individual mice. Graphs show mean±s.e.m. *p<0.05, ****p<0.0001. Group differences were tested using one-way ANOVA with Dunnett’s post-test (A) when variances were unequal or Kruskal–Wallis with Dunn’s post-test (B and C) for non-normally distributed data.

**Supp. Figure S3 | Central response to systemic inflammation in female BALB/c mice.** Brains were collected 1.5 h or 3 h post LPS or PBS (controls), sliced coronally, and imaged for Fos labeling and quantification. (**A-D**) From left to right: images from the Allen Mouse Brain Atlas, followed by coronal brain sections from PBS- and LPS-treated mice. Graphs show mean number of Fos+ cells per region per time point. (**E**) Summary of findings. Abbreviations: ARH, arcuate nucleus of the hypothalamus; pPVT, posterior portion of the paraventricular nucleus of the thalamus; CeA, central nucleus of the amygdala. Data are presented as mean±s.e.m. Two-way ANOVA was used to compare group differences. *p< 0.05, **p<0.01. Scale=170μm. Panel E was created with BioRender.

**Supp. Figure S5 | Peripheral LPS increases forced-swim immobility during the light phase.** Sickness behaviors were analyzed 24h post intraperitoneal injection of female BALB/c with 1 mg/kg LPS at ZT5. Control mice received PBS at the same time point. (**A**) Time spent in the center zone and (**B**) Time spent in the corners of the open field arena. (**C**) Distance traveled over 10 min in the open field arena. (**D**) Percentage of time immobile during the forced swim task. Graphs show mean±s.e.m. *p<0.05, ****p<0.0001. Group differences were tested using unpaired t-test (A and D) for normally distributed data or Welch’s t-test (C) when variances are unequal or Mann Whitney U test (B) for non-normally distributed data.

