## Supplemental Figure 1 for "Systemic endotoxemia induces integrated sickness physiology in female BALB/c mice"

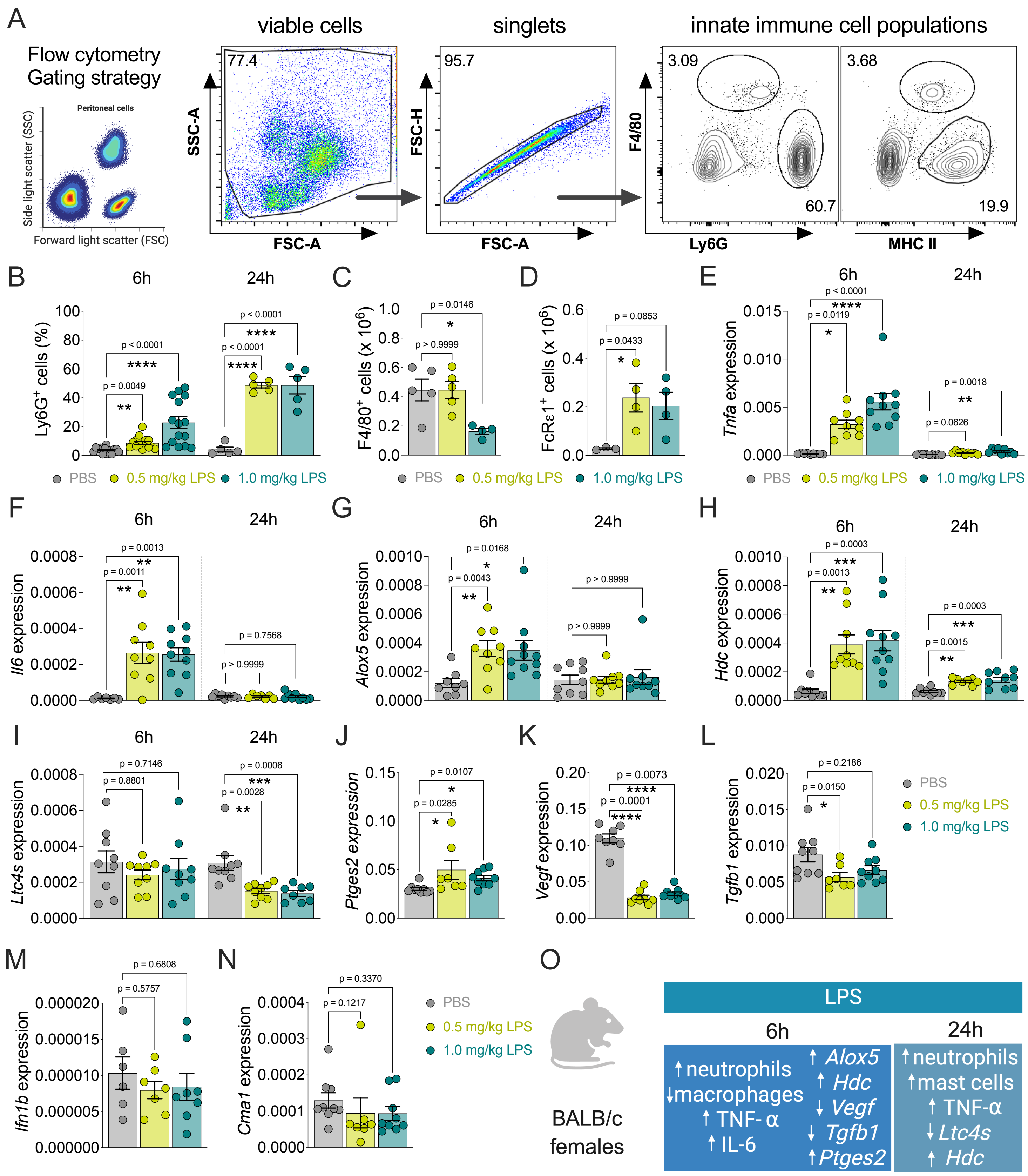

**Supp. Figure S1 | LPS treatment induces local and systemic changes in immune cells profile and gene expression in female BALB/c mice.** (**A**) Gating strategy used to determine the frequencies of innate immune cells in the peritoneal cavity of a high dose LPS-treated mouse by flow cytometry. (**B**) Percentage of Ly6G+ cells (neutrophils) in the peritoneal cavity of PBS- and LPS-treated mice at 6h and 24h post injection. (**C**) Number of F4/80$^{+}$ peritoneal cells 6h post LPS. (**D**) Number of FcεRI$^{+}$ peritoneal cells 24h post LPS. (**E-N**) Liver expression of tumor necrosis factor α (*Tnf*), interleukin 6 (*Il6)*, arachidonate 5-lipoxygenase (*Alox5)*, histidine decarboxylase (*Hdc*), leukotriene c4 synthase (*Ltc4s*), prostaglandin E synthase 2 *(Ptges2),* vascular endothelial growth factor (*Vegf),* transforming growth factor β1 (*Tgfb1),* interferon β 1 *(Ifnb1), and* chymase 1 (*Cma1)* relative to housekeeping gene *Rpl13a* quantified by RT-qPCR. **(J-N)** Panels show expression of liver transcripts at 6h post injections. (**O**) Summary of findings. Kruskal–Wallis test with Dunn's post-test (B-6h, C-24h, D, E, F-24h, G, H-6h, I-24h, J, K, L, N) or One-way ANOVA test with Dunnett's post-test (B-24h, H-24h, I-6h, M) was used. Data are presented as mean±s.e.m. *p<0.05, **p<0.01, ***p<0.001, ****p<0.0001. A and O were created with BioRender.
