## Supplemental Figure 2 for "Systemic endotoxemia induces integrated sickness physiology in female BALB/c mice"

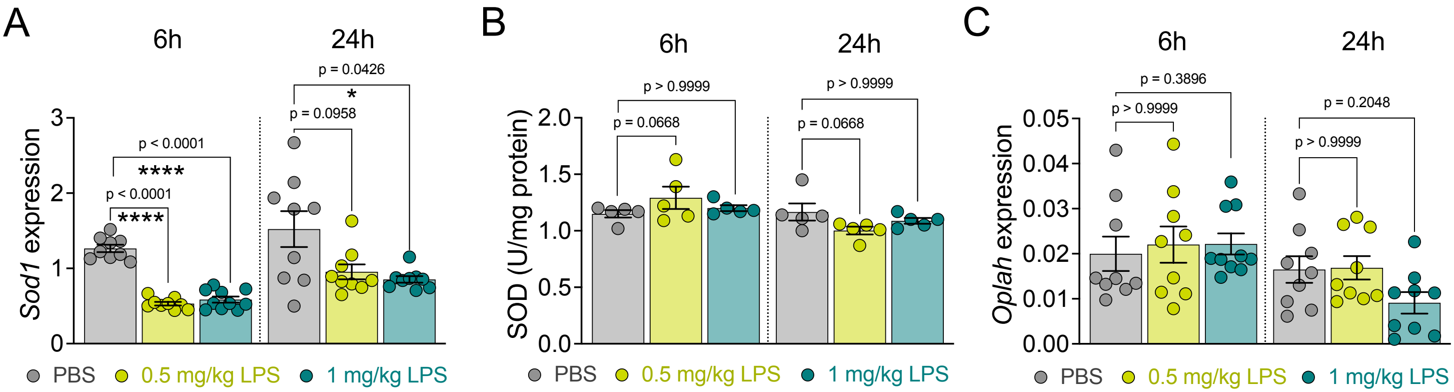


**Supp. Figure S2 | Hepatic antioxidant responses following LPS challenge. (A)** Hepatic expression of superoxide dismutase 1 (*Sod1*) at 6 h and 24 h after intraperitoneal injection of PBS, 0.5 mg/kg LPS, or 1.0 mg/kg LPS. **(B)** Superoxide dismutase (SOD) activity (U/mg protein) in the liver. **(C)** Hepatic expression of 5-oxoprolinase (*Oplah*) at 6 h and 24 h relative to housekeeping gene *Rpl13a* quantified by RT-qPCR. Points represent individual mice. Graphs show mean±s.e.m. *p<0.05, ****p<0.0001. Group differences were tested using one-way ANOVA with Dunnett's post-test (A) when variances were unequal or Kruskal–Wallis with Dunn's post-test (B and C) for non-normally distributed data.
