## Supplemental Figure 3 for "Systemic endotoxemia induces integrated sickness physiology in female BALB/c mice"

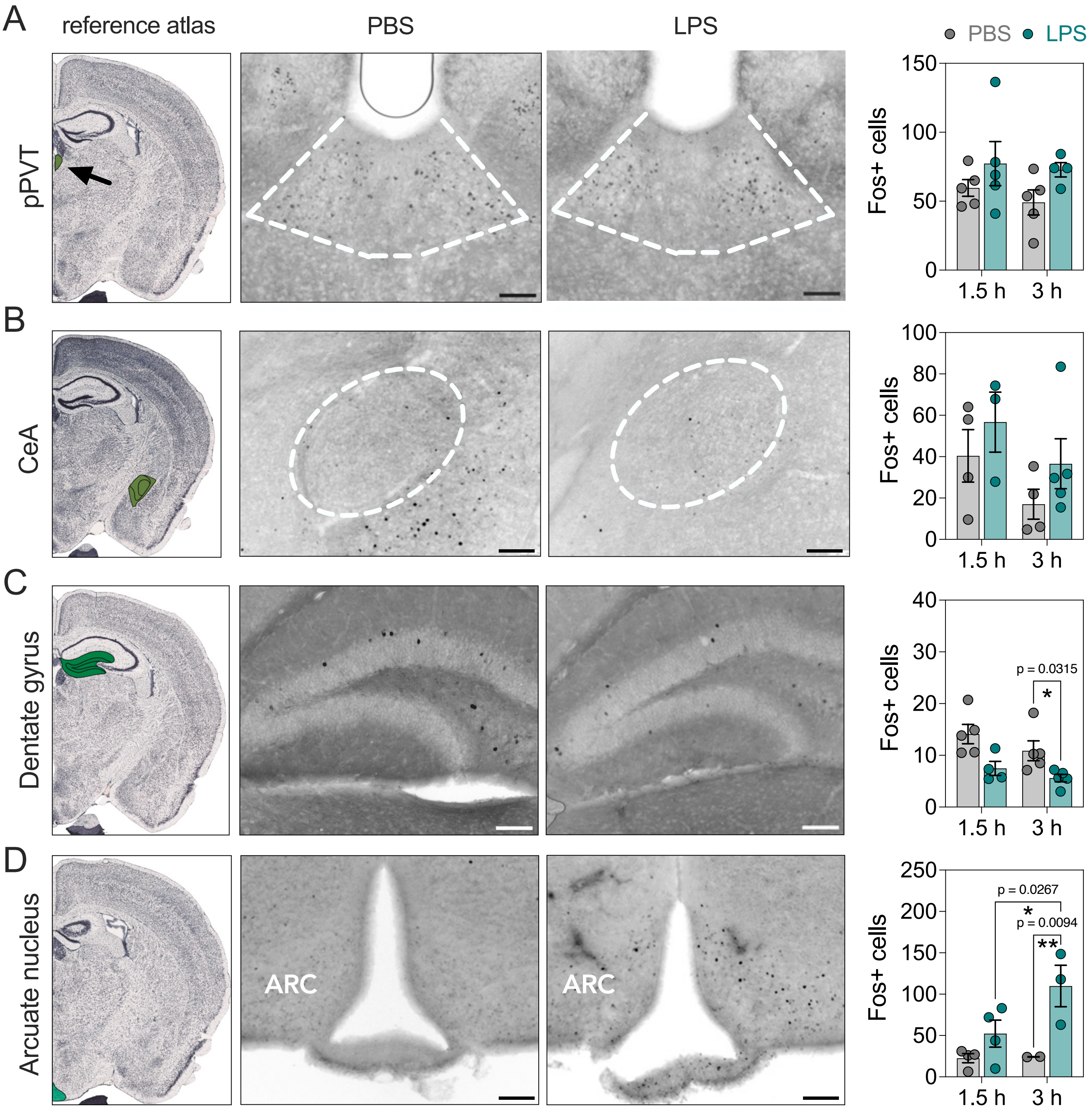


**Supp. Figure S3 | Central response to systemic inflammation in female BALB/c mice.** Brains were collected 1.5 h or 3 h post LPS or PBS (controls), sliced coronally, and imaged for Fos labeling and quantification. (**A-D**) From left to right: images from the Allen Mouse Brain Atlas, followed by coronal brain sections from PBS- and LPS-treated mice. Graphs show mean number of Fos+ cells per region per time point. Abbreviations: ARC, arcuate nucleus of the hypothalamus; pPVT, posterior portion of the paraventricular nucleus of the thalamus; CeA, central nucleus of the amygdala. Data are presented as mean ± s.e.m. Two-way ANOVA was used to compare group differences and time points. *p < 0.05, **p < 0.01. Scale = 170μm.
