## Supplemental Figure 4 for "Systemic endotoxemia induces integrated sickness physiology in female BALB/c mice"

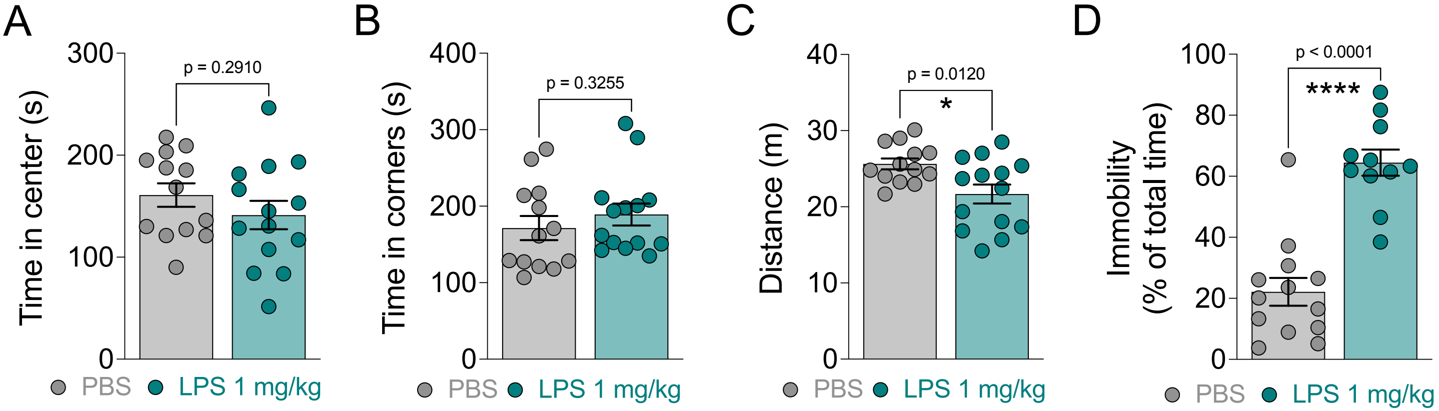


**Supp. Fig. 4 | Peripheral LPS increases forced-swim immobility during the light phase.** Sickness behaviors were analyzed 24h post intraperitoneal injection of female BALB/c with 1 mg/kg LPS at ZT5. Control mice received PBS at the same time point. (**A**) Time spent in the center zone and (**B**) time spent in the corners of the open field arena. (**C**) Distance traveled over 10 min in the open field arena. (**D**) Percentage of time immobile during the forced swim task. Graphs show mean±s.e.m. *$p<0.05$, ****$p<0.0001$. Group differences were tested using unpaired t-test (A and D) for normally distributed data or Welch's t-test (C) when variances are unequal or Mann Whitney U test (B) for non-normally distributed data.
